# Phosphorylation Promotes Liquid-Liquid Phase Separation of GRP8 and Its Assembly into Stress Granules Upon Salinity Stress in Arabidopsis

**DOI:** 10.1101/2023.12.13.571504

**Authors:** Adrian Kasztelan, Justyna Maszkowska, Anna Anielska-Mazur, Dominika Cieślak, Lidia Polkowska-Kowalczyk, Jarosław Poznański, Michał Dadlez, Christiane Nöh, Alexander Steffen, Karolina Kasztelan, Maria Bucholc, Katarzyna Patrycja Szymańska, Emilio Gutierrez-Beltran, Dorothee Staiger, Olga Sztatelman, Grażyna Dobrowolska

## Abstract

Drought and salinity are major environmental stresses affecting plant development and growth. SNF1–related protein kinases type 2 (SnRK2s) are key regulators of the plant responses to water deficit and salt stress. Here, we show that *Arabidopsis thaliana* Glycine-Rich RNA-Binding Protein 8 (GRP8) is a target of abscisic acid (ABA)-non-activated SnRK2s and negatively regulates root growth and seed germination under salt stress. In response to salinity, GRP8 assembles into stress granules (SGs). We show that in addition to the GRP8 C-terminal glycine-rich intrinsically disordered region (IRD), the N-terminal RNA recognition motif (RRM) plays a key role in this process. Phosphorylation of S27 in the RRM by SnRK2s significantly affects the structural dynamics of GRP8, facilitates its dimerization and subsequent liquid-liquid phase separation. Thus, we show that in addition to the known role of IDRs in recruitment into SGs, the RRM plays a decisive role.

## INTRODUCTION

Salinity is one of the most common stresses encountered by plants. Under salinity, seed germination, photosynthesis, growth, and metabolism are significantly inhibited. To survive, plants activate several signaling pathways controlling their defense mechanisms (for review, see ^1–3^). When the environment returns to optimal conditions, plants must switch back their metabolism from the stress response to growth. Some protein kinases, phosphatases, and RNA-binding proteins (RBPs) not only trigger and regulate stress responses, but also act as molecular switches between growth and stress defense. Recent studies have shown that RBPs can modulate plant development and sensitivity to environmental stresses by forming biomolecular condensates of different nature (for review, see ^4–6^).

The assembly of biomolecular condensates composed of a mixture of proteins (mainly nucleic acid-binding proteins), nucleic acids, and various small molecular weight molecules is driven by liquid-liquid phase separation (LLPS) (for review, see ^7–10^). The multivalent DNA/RNA-binding proteins with intrinsically disordered regions (IDRs) interacting with other molecules (proteins, RNA, or DNA) are core components of the condensates, where IDRs are usually the driving force behind LLPS. In this respect, animal and yeast RBPs, which have in their structure RNA binding motifs, e.g., RNA recognition motif (RRM), and IDR domains that lack a defined structure, e.g., rich in glycine interposed with aromatic and positively charged (very often arginines) amino acid residues have been intensively studied. It has been shown that weak electrostatic, hydrophobic, π–cation, and π–π interactions are responsible for LLPS (for review, see ^7,11^). In addition, protein post-translational modifications (PTMs), including phosphorylation, have been shown to play an important role in the regulation of LLPS and the assembly/disassembly of condensates ^12^; for recent reviews, see ^11,13–15^). To date, the information concerning the involvement of PTMs in regulating the formation and functioning of biomolecular condensates in plants is limited.

SNF1–related protein kinases type 2 (SnRK2s) are plant-specific protein kinases involved in the regulation of plant growth, development, and responses to various environmental stresses, especially osmotic stress, which in nature is a result of drought or salinity (for review, see: ^16,17^). All SnRK2s except SnRK2.9 in Arabidopsis are activated upon osmotic stress (for review, see ^16,17^). Based on their phylogeny, SnRK2s are divided into three groups (Extended Data Scheme 1). Our research focuses on SnRK2s from group 1, which comprises kinases that are strongly activated in response to osmotic stress but not to abscisic acid (ABA) (in Arabidopsis, SnRK2.1, SnRK2.4, SnRK2.5, and SnRK2.10). So far, data on their cellular targets are scarce.

Our earlier phosphoproteomic analysis indicated that several RBPs, glycine-rich RNA-binding protein 8 (GRP8) among them, could be targets of the ABA-non-activated SnRK2s ^18^. GRP8 comprises an N-terminal RNA recognition motif (RRM) domain and a C-terminal glycine-rich region. GRP8 and its paralog GRP7 are multifunctional RBPs involved in the post-transcriptional regulation of gene expression ^19–21^. Notably, an involvement of GRP7 in response to adverse environmental conditions has been reported ^22–24^. Considerably less research has been devoted to GRP8. Overall, GRP7 and GRP8 play similar roles in various processes, e.g., in the regulation of miRNA biogenesis ^21^ and alternative splicing ^20^, but their functions are not fully interchangeable ^25,26^.

Here, we investigated the role of GRP8 phosphorylation in the response to salinity. GRP8 is phosphorylated by the ABA-non-activated SnRK2s at serine 27 (S27), and this phosphorylation regulates its subcellular localization. Under regular growth conditions, GRP8 is localized to the nucleus and the cytoplasm, whereas upon salinity stress, most of the cytoplasmic GRP8 is translocated into stress granules (SGs). We show that the GRP8 assembly into SGs involves its phosphorylation at S27 in the RRM, which enhances the conformational dynamics of GRP8 and promotes its oligomerization and LLPS, eventually leading to its recruitment into SGs.

## RESULTS

### GRP8 is a Substrate of ABA-non-activated SnRK2s

An earlier study showed that in response to salinity stress, GRP8 was phosphorylated on S27 in Arabidopsis roots by SnRK2.10 or a kinase downstream of SnRK2.10 (^18^, Table S3). Now, using an *in vitro* phosphorylation assay, we showed that recombinantly expressed SnRK2.10 phosphorylates recombinant GRP8 (Extended Data Fig. 1). Additionally, the *in vitro* phosphorylation revealed that GRP8 could be phosphorylated not only by SnRK2.10 but also by other ABA-non-activated SnRK2s, SnRK2.4 or SnRK2.5 (Extended Data Fig.1). One of the phosphopeptides identified in GRP8 phosphorylated *in vitro* by SnRK2s (Extended Data Table 1) comprised phospho-S27 identified in the previous phosphoproteomic analysis. Furthermore, it was the primary phosphorylation site for ABA-non-activated SnRK2s (Extended Data Fig. 1). These results indicate unequivocally that SnRK2.10 and likely other ABA-non-activated SnRK2s phosphorylate GRP8 on S27 under salt stress conditions.

### GRP8 Negatively Regulates Germination and Root Growth under Salt Stress

The involvement of GRP8 in the response to salt stress has not been studied so far because no true *grp8* loss-of-function T-DNA line was available. To overcome this limitation, we used CRISPR-Cas9 and single guide RNAs 7 and 1 designed against regions in exon 1 and exon 2 and obtained *grp8* mutants with insertions/deletions (*grp8-5* and *grp8-11*) (Fig.1A, B). Immunoblotting with an anti-GRP8 antibody did not detect GRP8 protein in these lines, confirming a knockout (Fig. 1C). Notably, GRP7 was only slightly upregulated compared to the wild-type Col-0 (WT), indicating only a weak negative feedback from GRP8 on GRP7. This is contrary to the *grp7-1* mutant lacking GRP7, in which RNA and protein levels of GRP8 are elevated due to relief of repression by GRP7 ^19,27^.

**Figure 1.**
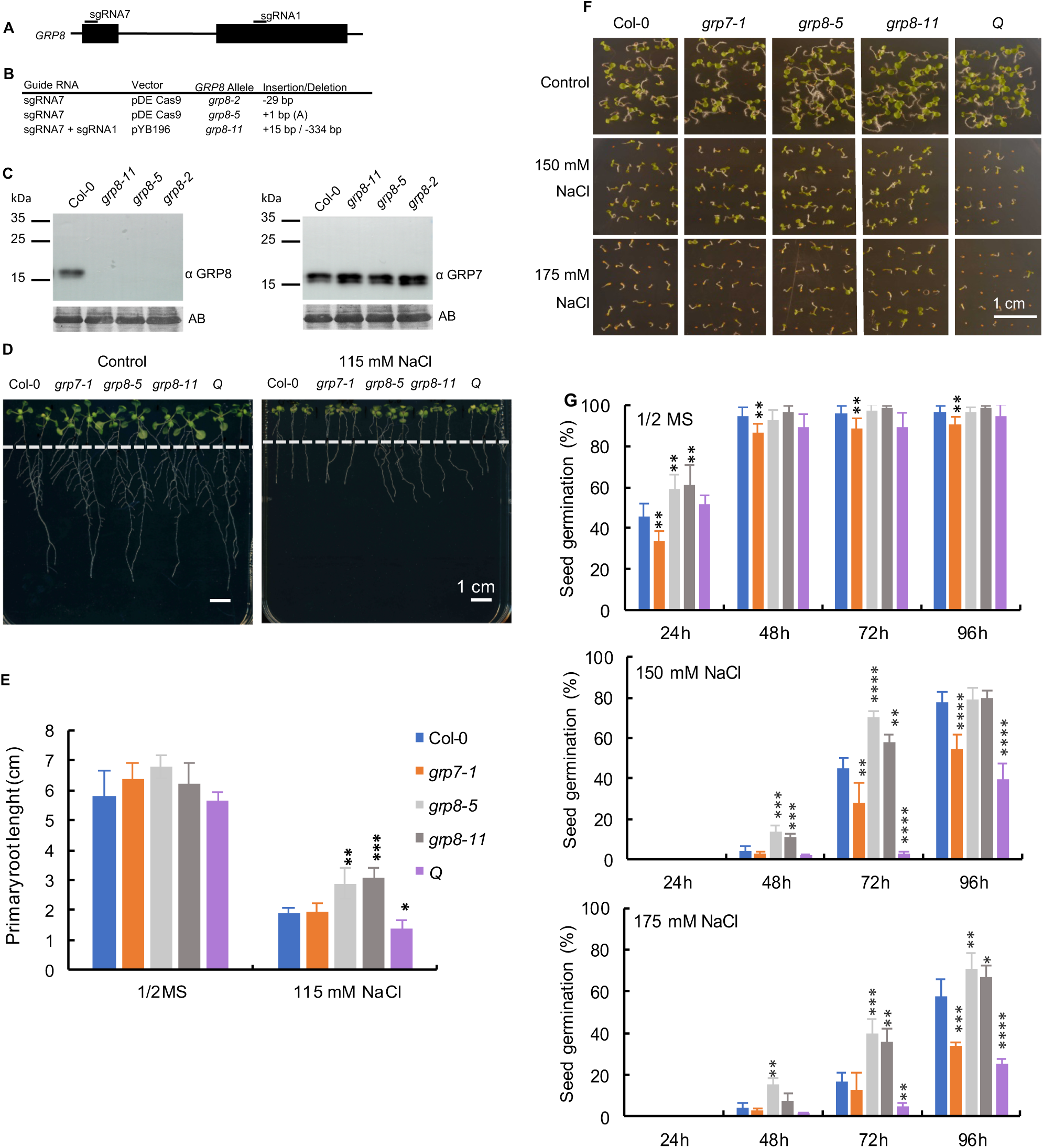
GRP8 regulates root growth and seed germination under salinity stress. (A) Schematic representation of the *AtGRP8* locus and positions of guide sequences used for CRISPR/Cas9 procedure (exons are shown as black boxes, and the intron is represented by a thin line); (B) Loss-of-function alleles of *GRP8* created with guide RNAs #1 and #7 and the nature of the mutations; (C) Expression of GRP8 and GRP7 proteins in wild type (Col-0) and *grp8* mutant Arabidopsis seedlings. For immunoblotting, anti-peptide antibodies against *At*GRP8 or *At*GRP7 were used. Amidoblack (AB) staining of the membrane served as a loading control; (D) Root growth phenotypes of wild type (Col-0), *grp7-1*, *grp8-5*, *grp8-11*, and *snrk2s* (Q) Arabidopsis mutants; (E) Quantification of the results from Figure 1D; (F) Seed germination phenotypes of the lines presented in Figures 1D and 1E at 7 d after sowing on ½ MS medium or medium supplemented with various concentrations of NaCl; (G) Quantification of the results from Figure 1F. Panels C, D, and F show representative results of three independent repeats of the experiment. The results presented in panels E and G are means with standard deviations. Statistical significance between groups was determined by One Way ANOVA and Student’s *t*-test. The asterisks indicate significant differences between WT Col-0 and mutants (*P <0.05, **P<0.01, ***P<0.001, ****P<0.0001).

Since salinity inhibits seed germination and root growth, we compared seed germination and primary root growth of the *grp8-5* and *grp8-11* knockout mutants, the *grp7-1* mutant, the *snrk2.1/2.4/2.5/2.10* quadruple mutant (Q) in which none of the group 1 SnRK2s is expressed ^18^, and WT Arabidopsis, in control and high-salt conditions (Fig. 1D, E, F, G).

Significant differences between the lines with respect to the primary root length were observed after salt stress application. The roots of the Q mutant were significantly shorter than the WT ones, which is consistent with previously obtained results for the similar *snrk2.1/2.4/2.5/2.10* mutant, which was generated by different alleles ^28^. In contrast, the roots of the two *grp8* mutants were significantly longer than those of all the other lines studied.

Additionally, an analysis of seed germination showed that the Q mutant was more strongly inhibited by salinity than were all the other lines studied, while the *grp8* mutants behaved just the opposite, showing a significantly higher germination rate than the other lines. Conversely, the *grp7-1* seeds germinated significantly slower than the WT or the *grp8* mutant seeds in both control and salinity stress conditions (Fig. 1F, G). The results indicate that SnRK2s and GRP8 play opposite roles in the response to salinity.

### Salinity Stress Triggers Phosphorylation-Dependent Recruitment of GRP8 into Stress Granules

Phosphorylation has been shown to trigger changes in the subcellular localization of numerous proteins, including RBPs ^29^. To monitor the subcellular localization of GRP8 under control conditions and upon salt stress, we examined roots of transgenic Arabidopsis plants stably expressing *pGRP8::msfGFP-GRP8* (msfGFP stands for <u>m</u>onomeric <u>s</u>uperfolder GFP) in the *grp8-11* background. In control conditions, GRP8 was evenly dispersed in the cytoplasm and the nucleus, while after salt stress application, when the SnRK2s are activated ^16^, GRP8 was present mainly in cytoplasmic foci (Fig. 2A and Extended Data Fig. 2). In parallel, for GRP8 localization we employed additional experimental models, *Nicotiana benthamiana* leaves transiently expressing *p35S::eGFP-GRP8* (Fig. 2B) and *Arabidopsis thaliana* plants stably expressing *p35S::eYFP-GRP8* (Fig. 3A). The GRP8 localization was the same in all models; in control conditions GRP8 was dispersed in the nucleus and the cytoplasm, while upon salt stress it was localized in cytoplasmic foci.

**Figure 2.**
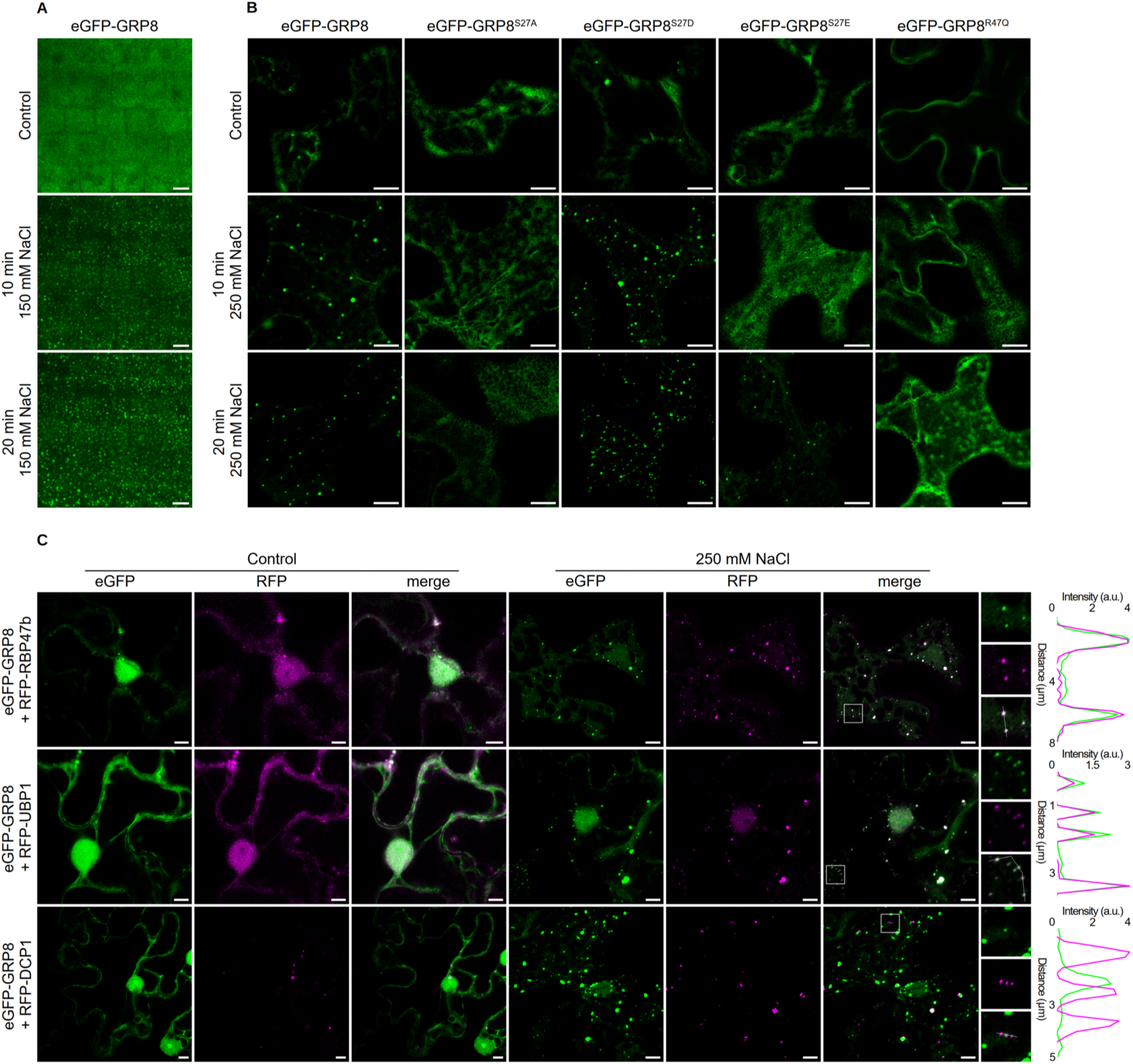
GRP8 is recruited into stress granules upon salt stress in a phosphorylation- and RNA binding-dependent manner. (A) Confocal images of msfGFP-GRP8 stably expressed from pGRP8::msfGFP-GRP8 in the roots of *grp8-11,* in control conditions and upon salt stress (B) Confocal images of eGFP-GRP8 and its variants in transiently transformed *N. benthamiana* epidermal cells exposed or not (control) to salt stress (250 mM NaCl). For interpretation of the GRP8 variants assayed, see text. (C) Co-localization of eGFP-GRP8 with RFP-tagged stress granule (UBP1b and RBP47b) and P-bodies (DCP1) markers in transiently transformed *N. benthamiana* epidermal cells, exposed or not to 250 mM NaCl for 20 min. Results of one of three independent experiments showing similar results are shown. Scale bar = 10 μm. Intensity scale values of co-localisation plots were divided by 1000.

**Figure 3.**
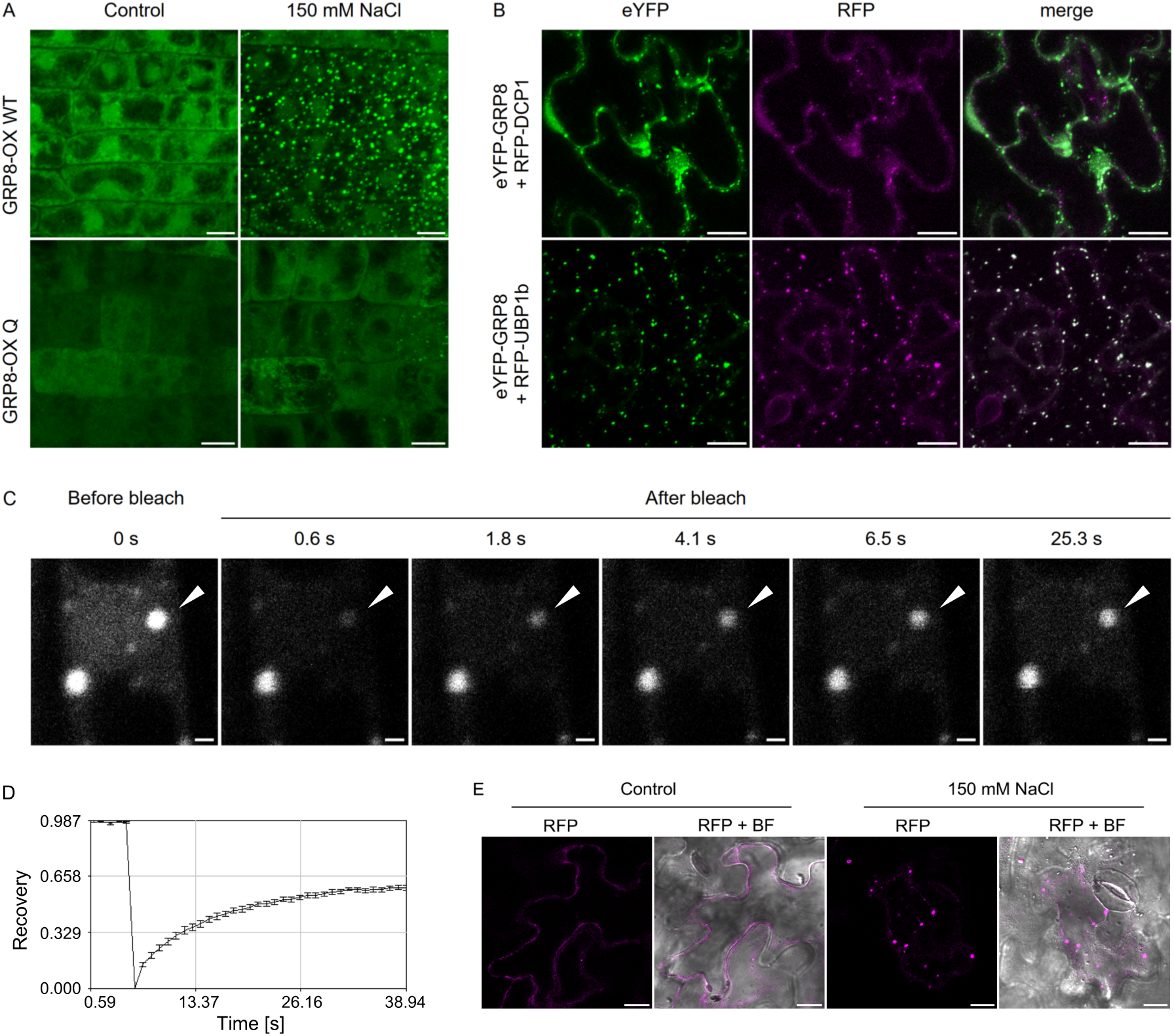
ABA-non-activated SnRK2s promote dynamic recruitment of GRP8 into stress granules. (A) Confocal images of eYFP-GRP8 in 7-day-old roots of eYFP-GRP8-WT and eYFP-GRP8-Q seedlings under control and salinity stress (150 mM NaCl, 60 min); (B) Co-localization of eYFP-GRP8 with RFP-UBP1b in eYFP-GRP8 WT leaves transiently transformed with a plasmid expressing RFP-UBP1b exposed to salinity stress (150 mM NaCl for 60 min). Results of one of three independent experiments showing similar results are shown. Scale bar = 10 μm; (C) FRAP of eYFP-GRP8 present in SGs formed in roots of eYFP-GRP8 WT seedlings subjected to salinity stress (150 mM NaCl for 60 min); bleached droplet indicated by an arrowhead; Scale bar = 1 μm (D) Time course of fluorescence recovery after photobleaching of eYFP-GRP8 present in SGs shown in Fig. 3A. Data are presented as the mean +/-s.d (n=10 independent experiments). (E) Confocal images of leaves of *grp8-11* mutant transiently expressing RFP-UBP1, exposed or not to salinity stress (150 mM NaCl for 60 min). Results of one of three independent experiments showing similar results are shown. BF = bright field image; Scale bar = 10 μm.

To check whether the S27 phosphorylation affects the GRP8 localization, we used the simplest approach, transient expression in *N. benthamiana.* We analyzed the subcellular localization of GRP8^WT^ and its non-phosphorylable (GRP8^S27A^) or phosphomimetic (GRP8^S27D^, GRP8^S27E^) variants in fusion with eGFP. All those GRP8 variants localized to the cytoplasm and the nucleus in control conditions. In contrast, GRP8^WT^ and its two phosphomimetic variants were additionally found in punctate structures after salt stress application. The non-phosphorylable GRP8^S27A^ variant failed to change its subcellular distribution upon salt stress (Fig. 2B). Interestingly, the GRP8^S27E^ variant showed a behavior in-between this of the GRP8^S27D^ and GRP8^S27A^ variants, indicating that glutamic acid inefficiently mimicked phosphoserine.

To determine whether the translocation of GRP8 requires its interaction with RNA, we analyzed the behavior of GRP8^R47Q^ with the conserved R47 in the RRM mutated, which markedly impaired its RNA-binding abilities ^19^; eGFP-GRP8^R47Q^ failed to exhibit a punctate distribution after salt stress (Fig. 2B), indicating that RNA binding is necessary for the GRP8 assembly into cytoplasmic foci.

The S27 residue is located within the RRM of GRP8; therefore, its phosphorylation could be expected to affect RNA binding. Our results showed that the S27 substitution with a phospho-mimicking residue (D) reduced GRP8 binding to an oligonucleotide derived from its own 3’ UTR, representing the best-known GRP8 target (described by ^19^, Extended Data Fig. 3). Therefore, while RNA binding seems to be necessary for translocation into foci, it is not enhanced by phosphorylation, leading to the conclusion that S27 phosphorylation is triggering the GRP8 translocation via another mechanism.

In eukaryotic cells, there are two major types of cytoplasmic foci where non-translating mRNAs, RBPs, and other proteins accumulate under adverse conditions: P-bodies (PBs) and stress granules (SGs) (for review, see ^4,30,31^). To determine whether the foci recruiting eGFP-GRP8 upon salt stress are related to either, we studied its co-localization with specific markers of SGs (RNA-binding protein 47b, RBP47b), and oligouridylate binding protein 1, UBP1) and PBs (decapping complex component 1, DCP1) fused with red fluorescence protein (RFP). We co-transformed *N. benthamiana* leaves with *eGFP-GRP8* and markers of SGs (*RFP*-*RBP47b* or *RFP*-*UBP1*) or PBs (*RFP*-*DCP1*); eGFP-GRP8 co-localized with RFP-RBP47b and RFP-UBP1 and not with RFP-DCP1 (Fig. 2C). These results indicated that following salt stress, GRP8 associates with SGs and additionally showed the commonly observed proximity of SGs and PBs ^30,32,33^.

To verify if, as inferred from the above data, the phosphorylation by the ABA-non-responsive SnRK2s promotes GRP8’s recruitment to SGs, we created transgenic plants expressing *p35S::eYFP-GRP8* in the wild-type (eYFP-GRP8-WT) and the *snrk2.1/2.4/2.5/2.10* mutant (eYFP-GRP8-Q) backgrounds. The levels of eYFP-GRP8 protein were similar in the eYFP-GRP8-Q and eYFP-GRP8-WT plants, and they were comparable to the levels of the native untagged GRP8 in non-transgenic plants (Extended Data Fig. 4A). Apparently, the ectopic expression of exogenous the GRP8 caused a slight decrease in the endogenous GRP8 level, demonstrating the correct negative autoregulation of GRP8 expression described earlier ^19^.

In roots of both types of transgenic plants, the cytoplasmic eYFP-GRP8 was dispersed under control conditions, while after NaCl treatment, most of it redistributed to cytoplasmic foci in the eYFP-GRP8-WT lines; this redistribution was much less pronounced in eYFP-GRP8-Q (Fig. 3A), indicating that GRP8 phosphorylation by SnRK2s promotes the relocalization. We confirmed the accumulation of eYFP-GRP8 in SGs upon salt stress through its co-localization with SG but not PB markers that were transiently expressed in Arabidopsis eYFP-GRP8-WT leaves (Fig. 3B). Notably, we showed a dynamic nature of the GRP8-containing granules by fluorescence recovery after photobleaching (FRAP) assay that demonstrated rapid exchange of eYFP-GRP8 in the granules with the surrounding cytoplasm (Fig. 3C, D). Moreover, the granule formation is transient (stress-correlated), as indicated by their dissolution after NaCl removal (Extended Data 4B). All the results described so far consistently show that GRP8 translocates to SGs upon salt stress and that this localization requires both the RNA binding by GRP8 and its phosphorylation at S27. *In vivo*, the ABA-non-activated SnRK2s carry out this phosphorylation following salt stress, thereby promoting the assembly of GRP8 into SGs. Furthermore, we showed that GRP8 is not indispensable for SG formation under salt stress since we observed the formation of SGs in the leaves of the *grp8-11* mutant (Fig. 3E).

### S27 Phosphorylation Increases Conformational Dynamics and Destabilizes the N-terminal part of GRP8

To decipher the mechanism by which S27 phosphorylation affects the ability of GRP8 to assemble into SGs, we performed homology modeling of its N-terminal RRM phosphorylated or non-phosphorylated at S27. The best model of the GRP8 RRM was one based on a combination of 4CQ7 and 5TBX pdb structures of homologous domains deposed in the Protein Data Bank, while no reasonable templates were identified for the remaining C-terminal glycine-rich part of the protein. The hydroxyl group of S27, located at the C-terminus of the helix α1 (residues T17-S27), forms a hydrogen bond with the Q23 sidechain carbonyl, thus stabilizing the helix. Notably, phosphorylation of S27 breaks this interaction. Moreover, an unfavorable electrostatic interaction between pS27 and D31 side-chain carboxylic group decreases the structural stability of the βαββαβ fold (Fig. 4A). Overall, the phosphorylation of S27 was assessed with FoldX5 to destabilize RRM by 1.3 ± 0.2 kcal/mol. A similar effect of 1.1 kcal/mol was associated with the S27D replacement, while S27A and S27E only minutely affected the protein stability (< 0.15 kcal/mol).

**Figure 4.**
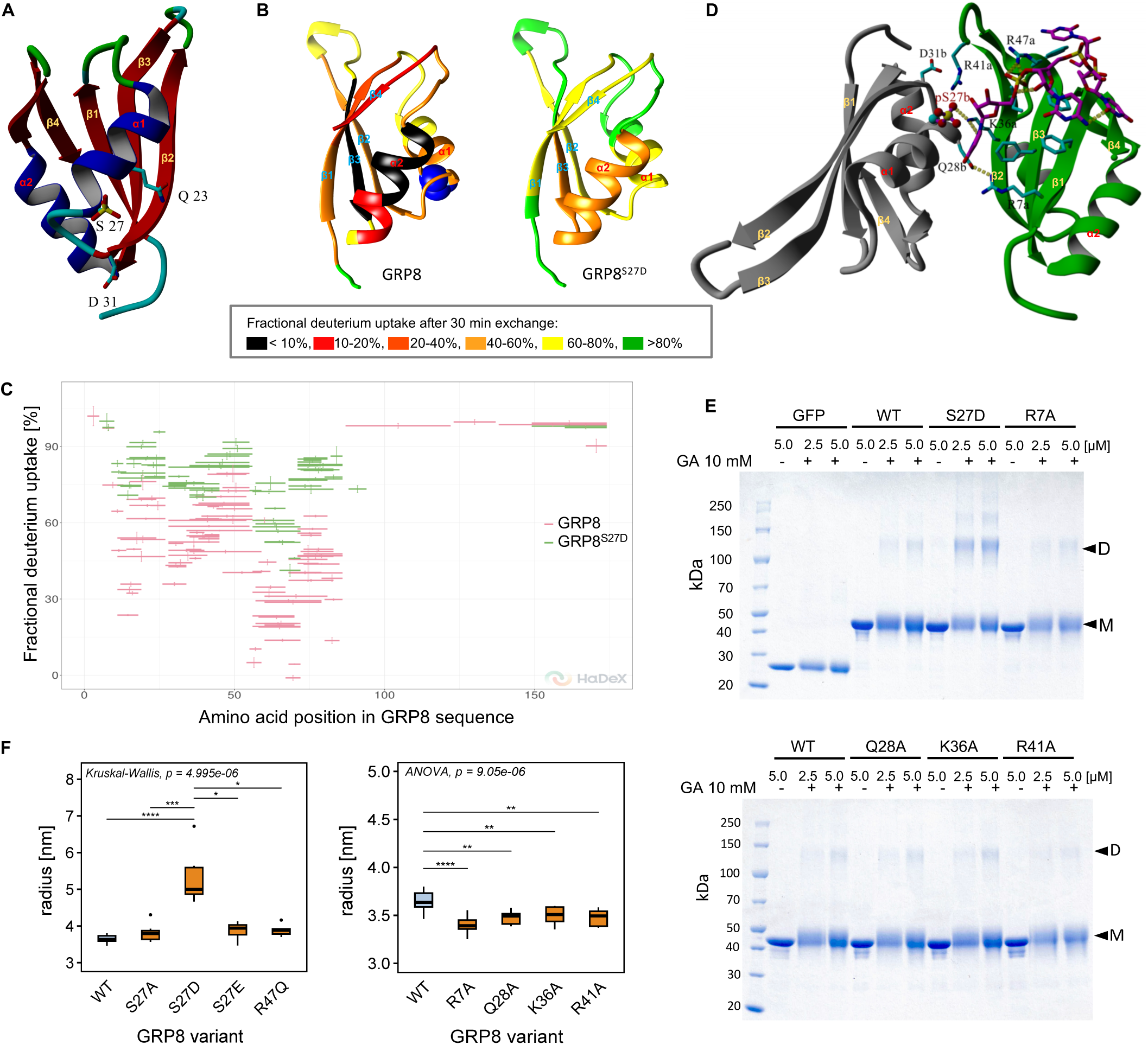
S27 phosphorylation affects protein conformational dynamics and dimerization of GRP8. (A) Model of GRP8 N-domain with pS27, Q23 and D31 represented as sticks; (B) Fractional deuterium uptake after 30 minutes of exposition to D2O for different regions of GRP8^WT^ and GRP8^S27D^ overlaid on the structural model of the N-terminal GRP8 domain. The exchange level is color-coded as described in the inset. The blue balls in GRP8^WT^ mark S27 position; (C) Extent of deuterium uptake by GRP8^WT^ and GRP8^S27D^ after 30 min of incubation with D2O along the entire GRP8 sequence; (D) Model of GRP8-GRP8 dimer with RNA, RNA represented as magenta sticks, amino acid residues crucial for dimerization as cyan sticks; (E) SDS-PAGE analysis of dimerization of GFP-GRP8^WT^ and GFP-GRP8 variants after glutaraldehyde (GA) crosslinking. The analyzed variants, except the phosphomimetic S27D, carried mutations in positions predicted to affect dimer stability. GFP alone was run as a negative control; M –GFP-GRP8 monomer, D – dimer; (F) Hydrodynamic radius (RH) of GFP-GRP8 variants determined by DLS. Results of one of ten independent experiments showing similar results are shown. Statistical significance of differences between variants using Kruskal-Wallis test with Dunn’s post-hoc comparison and Bonferroni p-value correction or one-way ANOVA with post-hoc Tukey’s HSD test (as indicated). *P <0.05, **P<0.01, ***P<0.001, ****P<0.0001. In Fig. 4C, 4D, and 4E, results of one of three independent experiments showing similar results are shown.

To verify experimentally the predicted destabilizing effect of S27 phosphorylation on the structure of GRP8, we compared the conformational dynamics of GRP8^WT^ and its phosphomimetic form GRP8^S27D^ using hydrogen/deuterium exchange monitored by mass spectrometry (HDX-MS). Figures 4B and 4C show the extent of fractional deuterium uptake in different regions GRP8^WT^ and GRP8^S27D^ of the folded N-terminal RRM subdomain (residues 1-80) superimposed on the GRP8 N-terminal part model (Fig. 4B) and along the entire GRP8 sequence (Fig. 4C) after 30 minutes of incubation. For both GRP8 variants, only its N-terminal RRM subdomain shows some degree of limitation of deuterium uptake, consistent with a folded structure; the C-terminal part is completely unstructured, as evidenced by its free hydrogen/deuterium exchange. For the folded part, the exchange rate is much slower for the WT than for the phosphomimetic variant S27D. For WT (Fig. 4B), significant protection is restricted to a region comprising the beta-strand β3 at positions 49-54 and the following helix α2 (residues 57-67) (marked red and black in the figure). For the loop between the β2 and β3 strands (residues 41-48) and the C-terminal IDR (residues 82-168), including the glycine-rich region, no significant protection against exchange was observed. In GRP8^S27D^ (Fig. 4B and C), β3 strand and α2 helix (51-67) are the only regions retaining some protection. Such profound differences in HDX between GRP8^WT^ and GRP8^S27D^ were observed at all incubation times, as illustrated by the deuterium uptake curves for selected peptides covering different regions of GRP8 (Extended Data Fig. 5A). Fitting the deuterium uptake curves to an exponential model indicated an overall 10-fold increase in the protein structure opening rate, corresponding well with the >1 kcal/mol destabilization of the GRP8 structure assessed in computer modeling, both for phosphorylated WT (pS27) and the GRP8^S27D^ variants. The HDX-MS experiments convincingly confirmed the earlier predictions from modeling by showing that S27D substitution and, by inference, the S27 phosphorylation markedly destabilizes the entire N-terminal domain of GRP8.

### S27 Phosphorylation Promotes GRP8 Dimerization

The GRP8 modeling additionally demonstrated that the RRM domain of GRP8 molecules can form an asymmetric dimer (Fig. 4D) which could further form higher-order quasi-polymeric structures (Extended Data Fig. 5A). S27 is located at the intermolecular interface of the putative GRP8 dimer, its phosphorylation should stabilize the dimer by a favorable short-range intramolecular electrostatic interaction with R41 (Fig. 4B), thus affecting RNA (denoted in magenta in Fig. 4D) binding. The dimer stabilization caused by S27 phosphorylation was estimated at -0.5 kcal/mol; the S27D replacement had a similar stabilizing effect (-0.6 kcal/mol), and S27A was neutral.

In the final step, the ternary complex of the GRP8 homodimer in complex with RNA was built and found to be stable in terms of 5 ns unconstrained Molecular Dynamics. Inspection of the resulting structure pointed to several interactions possibly involved in dimerization and RNA binding. The interdomain interface is stabilized by salt bridges formed by sidechains of R7(mol a):Q28(mol b) (∼1.5 ± 1.0 kcal/mol decrease for either R7A or Q28A replacements), K36(a):pS27(b) (∼0.5 ± 0.4 kcal/mol for either K36A or pS27A), and R41(a):D31(b) (decrease ∼1.7 ± 0.7 kcal/mol for R41A). Critical for the RNA binding was R47(a), whose guanidinium group interacted with three phosphate groups, those of U_i+2_, U_i+3_, and U_i+5_ (Fig. 4B).

To verify experimentally the ability of GRP8 to dimerize and form larger assemblies, we cross-linked GRP8 in solution using glutaraldehyde and examined the resulting GRP8 forms by SDS-PAGE. To ensure efficient cross-linking, GRP8 or its variants were fused msfGFP (further referred to as GFP) because, according to our model, the four lysine residues of GRP8 (K36, K55 K58, K70) are in an unfavorable orientation precluding glutaraldehyde crosslinking. The results revealed that, indeed, GRP8 is capable of forming dimers in solution and that GRP8^S27D^ forms dimers and higher-order complexes more efficiently than GRP8^WT^ (Fig. 4E). GRP8 variants mutated in residues predicted to participate in the dimerization showed only a slightly decreased abundance of the dimer; the effect was the most pronounced for GRP8^R7A^ and GRP8^R47A^ variants.

The above analysis was extended with a dynamic light scattering study, which showed that the hydrodynamic radius (R_H_) of GFP-GRP8^S27D^ was significantly larger than that of GFP-GRP8^WT^ or the other GRP8 variants, indicating that a significant fraction of GRP8^S27D^ exists as a dimer or higher-order oligomers in solution (Fig. 4F). The mutants designed to impact the dimerization had a slightly lower R_H_ than GRP8^WT^, suggesting a more compact conformation.

### GRP8 Undergoes Liquid-Liquid Phase Separation *In Vitro*

GRP8 contains a single RRM at the N-terminus (where S27 is located) and an IDR at the C-terminus. The C-terminal part is rich in glycine stretches interspersed with serine, tyrosine, and arginine residues and is predicted with a high probability to be a prion-like domain (PLD) (Fig. 5A), suggesting that GRP8 can undergo LLPS. To verify this hypothesis and to establish the respective roles of the N- and C-terminal domains of GRP8 in the predicted phase separation, we performed an *in vitro* LLPS assay. The following recombinant proteins were studied: GFP-GRP8^WT^ (full-length GRP8), GFP-N-GRP8 (N-terminal domain of GRP8, residues 1-82), and GFP-C-GRP8 (C-terminal half of GRP8, residues 83-169), and GFP alone as a negative control. In optimized LLPS assay conditions (Extended Data Fig. 6A and B, 12% PEG; 125 mM KCl in 25 mM HEPES-NaOH, pH 7.5), GFP-GRP8^WT^ underwent phase separation at a concentration of 20 μM (Fig. 5B), which was abolished upon addition of 1,6-hexanediol (Extended data Fig. 6F). Droplets coalescence and the FRAP analysis confirmed the dynamic nature of the *in vitro* formed condensates (Extended data Fig. 6D, G and H).

**Figure 5.**
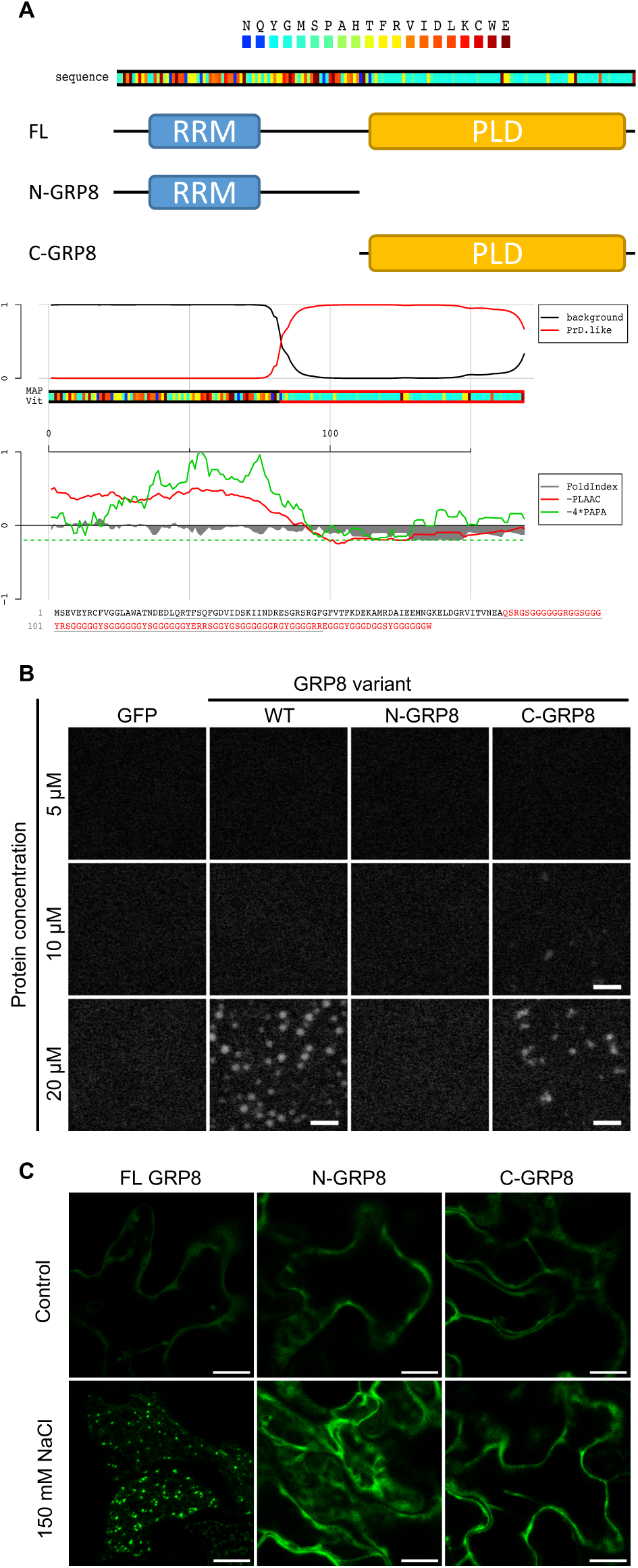
GRP8 contains prion-like domain and undergoes LLPS *in vitro*. (A) Amino acids distribution along GRP8 sequence, schematic representation of GRP8 truncations and PLD (prion-like domain) prediction; (B) *In vitro* LLPS of msfGFP-GRP8 FL and its truncated forms in the presence of PEG 3350; (C) *In vivo* LLPS by full-length (FL) GRP8 and its N- and C-domains fused N-terminally with eGFP in transiently transformed leaves of *grp8-11 Arabidopsis*;

GFP-C-GRP8 formed droplets less efficiently compared with GFP-GRP8^WT^, and their morphology was altered, whereas the GFP-N-GRP8 lacking the IDR region failed to form condensates at all concentrations tested (Fig. 5B). These results showed an absolute requirement for the C-terminal unstructured fragment of GRP8 for LLPS, but also suggested an involvement of the N-terminal domain in the transition. Such a cooperative mechanism of both domains was confirmed *in planta*; neither eGFP-N-GRP8 nor eGFP-C-GRP8 transiently expressed in the *grp8-11* mutant leaves was recruited to cytoplasmic foci upon salt stress (Fig. 5C).

The effect of S27 phosphorylation on GRP8 LLPS was examined using GFP-GRP8^WT^ and a set of phosphosite-GRP8 variants (GFP-GRP8^S27D^, GFP-GRP8^S27E^, GFP-GRP8^S27A^). At the highest protein concentration assayed (20 μM), all these GRP8 variants formed droplets, which were most prominent in the case of GFP-GRP^S27D^, and the fraction of phase-separated protein was the highest (Fig. 6A). Notably, at lower protein concentrations, only the S27D variant consistently exhibited efficient LLPS, indicating that this phosphomimetic mutation significantly lowered the critical protein concentration required for phase separation. One is tempted to assume that *in vivo* S27-phosphorylated GRP8 would also undergo phase separation more readily.

**Figure 6.**
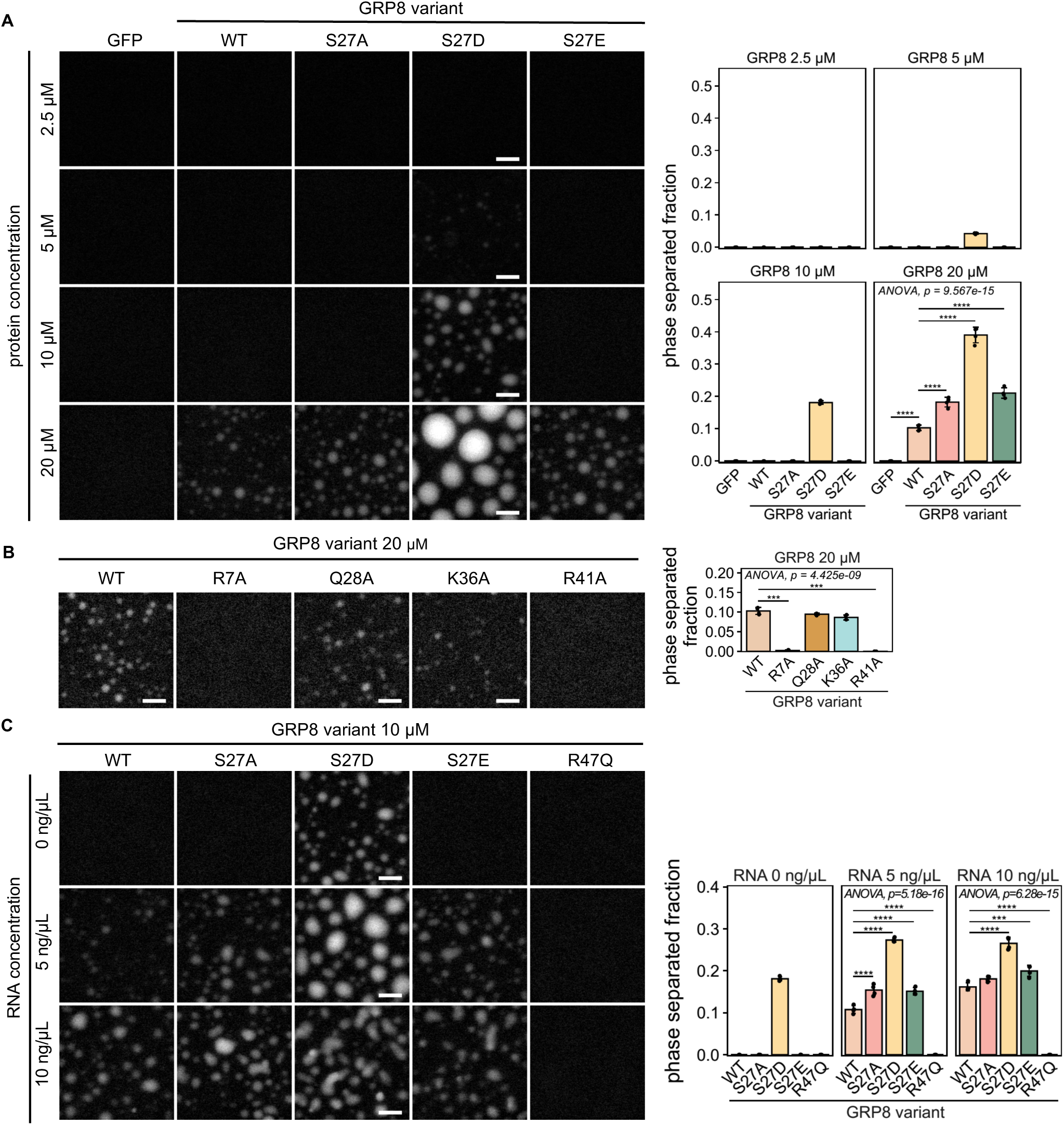
*In vitro* LLPS of GFP-GRP8 and its variants. (A) In vitro LLPS of purified msfGFP-GRP8 and its variants; Statistical significance of differences between variants was determined using one-way ANOVA with Dunnett’s post-hoc comparison. *P <0.05, **P<0.01, ***P<0.001, ****P<0.0001. (B) LLPS of GRP8 putative dimerization mutants; Statistical significance of differences between variants was determined using one-way Welch’s with Games-Howell ANOVA with Dunnett’s post-hoc comparison. *P <0.05, **P<0.01, ***P<0.001, ****P<0.0001. (C) Effect of total Arabidopsis RNA addition on LLPS of msfGFP-GRP8 and its variants; Statistical significance of differences between variants was determined using one-way ANOVA with Dunnett’s post-hoc comparison. *P <0.05, **P<0.01, ***P<0.001, ****P<0.0001. All assays depicted were carried out in the optimized LLPS buffer(12% PEG; 125 mM KCl in 25 mM HEPES-NaOH, pH 7.5). Results of one of three independent experiments showing similar results are shown. Scale bar = 10 μm.

Additionally, we tested *in vitro* LLPS for GFP-GRP8 variants, predicted to be affected in their dimerization ability: GFP-GRP8^R7A^, GFP-GRP8^Q28A^, GFP-GRP8^Q36A^, and GFP-GRP8^R41A^. GRP8^R41A^ and GRP8^R7A^ did not form droplets or formed only very few, respectively, even at the highest concentration tested (20 μM), whereas GRP8^Q28A^ and GRP8^Q36A^ exhibited only a slightly lower tendency to undergo LLPS than GRP8^WT^ (Fig. 6B). This lack of reciprocality suggests that during formation of GRP8 oligomers R7 and R41 may be involved in interactions with more amino acid residues than predicted from *in silico* modeling data. Nevertheless, substituting R7 or R41 with alanine severely disrupts LLPS, indicating that those residues are crucial for providing intermolecular interactions during phase separation.

### GRP8 – RNA Interaction Facilitates LLPS

It has been shown for a number of RBPs that their association with RNA greatly facilitates their LLPS ^10,34,35^. To determine if this is also the case for GRP8, we first checked its ability to undergo LLPS in a PEG-free solution but supplemented with total RNA from Arabidopsis seedlings; no LLPS was observed (Extended Data Fig. 6C). Next, we monitored the effect of RNA addition on the efficiency of phase separation of various GRP8 variants in the optimal conditions, i.e., in the presence of 12% PEG. GFP-GRP8^WT^, GRP8^S27D^, GRP8^S27E^, GRP8^S27A^, and additionally GFP-GRP8^R47Q^, with a significantly reduced RNA affinity, were assayed at a constant protein concentration of 10 μM. In these conditions, we observed droplet formation for GRP8^WT^, GRP8^S27E^, and GRP8^S27A^, which, crucially, did not undergo LLPS at 10 μM without the addition of RNA (Fig. 6C). RNA also enhanced the efficiency of the LLPS of GRP8^S27D^. In contrast to all other GRP8 variants, GRP8^R47Q^ at 10 μM did not undergo LLPS even in the presence of RNA, whereas at 20 μM (and in the absence of RNA), it did so (Extended Data Fig. 6E). Thus, enhancing the LLPS in the presence of RNA is strictly linked with RNA-protein interactions and not just presence of another macromolecule acting as a molecular crowding agent.

The above results show that oligomerization and the interaction with RNA significantly facilitate the LLPS of GRP8. The phosphorylation at S27 decreases the critical concentration of GRP8 LLPS by promoting GRP8 oligomerization.

## DISCUSSION

In response to salinity, ABA-non-activated SnRK2s phosphorylate GRP8 at S27 localized in the RRM, the phosphorylation of which promotes the recruitment of GRP8 into SGs. The mechanisms of SG formation and dissolution, as well as their exact role, are poorly understood ^36^, albeit numerous reports point to their function in the sequestration, stabilization and storing of mRNAs, RBPs, and other translation-related proteins (for review, see: ^9,30,31^). This allows translation to resume as soon as the stress is over and the conditions become suitable for growth again. Notably, diverse proteins and RNA species involved in stress signaling can be found in SGs, indicating that these condensates may also play a role in recruiting and bringing together components engaged in the stress response to increase the effectiveness of the defense or to exclude those interfering with it ^37–40^. ^40^ have shown that LLPS of two glycine-rich RNA-binding proteins, RBGD2 and RBGD4, and their assembly into SGs protect plant cells against heat stress. Also, ALBA proteins, which separate into SGs and PBs upon heat stress, interacting with and stabilizing *HSF* mRNAs, play a protective role under stress ^39^.

The recruitment of proteins to SGs, their assembly, fusion, and finally disassembly are tightly controlled. However, the knowledge about the signaling pathways responsible for the modulation of these processes remains scant, especially in plants. Recent studies on mammalian models show that post-translational modifications, including phosphorylation, affect the phenomenon of LLPS responsible for the formation of SGs, thereby regulating their assembly and disassembly (for review, see ^11,15^). To date, most data have shown a role of phosphorylation within the low-complexity/IDR of RBPs ^12,41,42^, whereas our results show that SnRK2s phosphorylate GRP8 at S27 located in the well-folded RMM domain. Quite unexpectedly, this phosphorylation promotes GRP8 assembly into SGs.

To underpin the mechanism of the GRP8 assembly into SGs, we analyzed *in silico* and experimentally several features of GRP8 and its mutated variants. Molecular modeling of the N-terminal RRM of GRP8 predicted that both phosphorylation of S27 and the phosphomimetic mutation S27D should destabilize α1 helix within the RRM domain while stabilizing the GRP8-GRP8 dimer/oligomer formation. These predictions were fully confirmed using biochemical and biophysical methods. Crucially, an *in vitro* LLPS assay showed that phosphorylation at S27 enhanced the GRP8 propensity to undergo phase separation.

The mechanism of SG formation in animal cells presented by Guillen-Boixtet (2020) has revealed that Ras-GTPase-activating protein (SH3 domain)-binding protein G3BP, which drives the SG assembly, relies on LLPS, which is triggered by changes in its conformational dynamics, on the ability of G3BP to dimerize, and on multiple protein-protein and protein-RNA interactions. Based on our present results, we propose a somewhat similar mechanism of GRP8 assembly into SGs in plant cells, with the caveat that, unlike G3BP, GRP8 is not crucial for SG formation. In response to salinity stress, SnRK2s phosphorylate a pool of GRP8 at S27; the phosphorylation increases the structural dynamics of GRP8 and modifies its intramolecular interaction network, which in turn facilitates interactions with other GRP8 molecules, eventually causing LPPS. Interactions with some proteins and RNAs may be altered, leading to GRP8 sequestration into the proper (SG) condensate; however, the latter needs further studies. While our data show that RNA binding is required for the GRP8 assembly into granules, a negative effect of the S27 phosphorylation on the RNA-GRP8 binding seems contradictory to this model. However, it should be noted that we analyzed the binding of short oligoribonucleotides, whereas SGs contain long unfolded RNA species, mainly mRNAs released from polysomes under stress conditions ^35^.

Our results indicate that GRP8 is phosphorylated by SnRK2s upon salt stress and regulates plant responses to salinity at the seed germination and root growth level. Notably, we found that the action of GRP8 is opposite to that of the SnRK2s in these processes.

A role of the ABA-non-activated SnRK2s in the regulation of root growth and architecture in response to salinity and drought has been shown before ^28,43,44^ and our results are in line with those studies. ^28^ and ^44^ linked the effect of the ABA-non-activated SnRK2s on root growth with the regulation of mRNA decay by these kinases through phosphorylation of VARICOSE (localized in PBs). ^44^ additionally suggested that the ABA-non-activated SnRK2s impact the root growth and architecture via another mechanism - by regulating the expression of genes encoding aquaporins PIP2;3 and PIP2;5, and the CYP79B2 involved in auxin biosynthesis. The present results indicate yet another scenario: upon salinity, the ABA-non-activated SnRK2s phosphorylate and trigger LLPS of GRP8, a negative regulator of PR growth and seed germination under salinity stress conditions, leading to its sequestration in SGs. Further studies should determine the network of the GRP8 protein-protein and protein-RNA interactions under salinity stress and define the role of its recruitment into SGs.

Concluding, we showed that GRP8 is phosphorylated at S27 by the ABA-non-activated SnRK2s in response to salinity. Upon stress, GRP8 is recruited to the SGs, which is promoted by the S27 phosphorylation, increasing the structural dynamics of GRP8 and its tendency to dimerize/oligomerize and undergo LLPS.

## MATERIALS AND METHODS

### Plant material, growth conditions, and stress treatment

Arabidopsis (*Arabidopsis thaliana*) ecotype Columbia 0 (Col-0) wild type; mutants: quadruple T-DNA insertion knockout mutant Q: *snrk2.1/2.4/2.5/2.10* (SAIL_519_C01/Salk_080588/Salk_075624/WiscDsLox233E9) described by ^18^; *grp7-1* mutant described by ^27^. Seeds were germinated, and plants were grown under long-day conditions (16 h light/8 h dark photoperiod) at 22°C/19°C on soil, in hydroponic culture, or on ½ MS (Duchefa) agar medium in Petri plates. For aseptic cultures, seeds were sterilized in 70% ethanol for 2 min, then in 10% (v/v) sodium hypochlorite solution for 20 min. After sterilization, the seeds were washed extensively with sterile water. Seeds were stratified in the dark at 4°C for three days. For transient expression experiments, *Nicotiana benthamiana* plants were grown in soil in a growth chamber under 60% relative humidity, at 16 h light/8 h dark, 23°C/19°C, while Arabidopsis plants were grown identically but in short day conditions (8 h light/16 h dark).

Arabidopsis root growth assay was performed as described in ^43^. Root length was determined using the ImageJ software (http://imagej.nih.gov/ij/). Differences in primary root length were determined using Microsoft Excel 2016 by One Way ANOVA and Student’s t-test.

Seed germination assay was performed basically as described by ^45^ with NaCl added to the medium instead of ABA. Briefly, sterilized seeds were sown on ½ MS medium with 10 g/L agar supplemented (or not) with NaCl at a concentration of 150 or 175 mM and stratified at 4°C for three days in darkness. Germination (emergence of radicles) was scored for up to 4 days after transferring plates into 16 h light/8 h dark photoperiod at 22/20°C. Differences in the germination rate were determined using Microsoft Excel 2016 by One Way ANOVA and Student’s t-test.

### Generation of *grp8* mutants via CRISPR/Cas9

The guide sequences ACGTGTCATCACCGTGAACG (#1) and GTTGAGTACCGGTGCTTTGT (#7) specific for *GRP8* were selected using CRISPR Design and Breaking–Cas ^46,47^. Both strands were synthesized as oligonucleotides flanked with BbsI restriction sites, annealed, and subcloned into pEN-C1.1 ^48^, providing the U6-26 promoter and the sgRNA scaffold sequence. The assembled sgRNAs were cloned into the binary vector pDe Cas9 ^48^ in a gateway LR reaction using LR clonase II (ThermoFisherScientific, USA) according to the manufacturer’s instructions.

To target the CRISPR constructs selectively to dividing tissues and obtain larger deletions, both sgRNAs were cloned into pYB196, providing Cas9 under the control of the *ICU2* promoter ^49^. The sgRNA #1 cassette was amplified from pEN-C1.1 with the primers F_pEN_NotI and R_pEN_NotI and ligated into pYB196. sgRNA #7 was amplified with the primers F_BamHI_pEN and R_SpeI_pEN and ligated into pYB196 already containing sgRNA #1. pDE Cas9 and pYB196 were introduced into agrobacterium strain GV3101 carrying pSOUP, and plants were transformed using the floral dip method.

Primary transformants were tested for the presence of the Cas9 constructs by spraying soil-grown plants with BASTA® solution. The activity of Cas9 at the *GRP8* locus was checked by employing a heteroduplex assay, essentially as described by^50^. Briefly, *GRP8* was amplified with primers CCRB_5’UTR and AGRP95, the PCR fragments were heated to 95°C and cooled to RT for re-annealing, and separated on native 15% polyacrylamide TBE gels. The formation of a heteroduplex indicated the presence of mutated *grp8* allele in a heterozygous state. In the following generations, homozygous mutant plants were identified by mixing *grp8* amplicons from putative mutants with non-mutated WT *GRP8* amplicons to allow heteroduplex formation. The homozygous *grp8* mutants were characterized in the T3 generation by Sanger sequencing and checked for the absence of GRP8 and the level of GRP7 with anti-peptide antibodies directed against GRP8 and GRP7. The absence of the Cas9 construct was tested by painting one leaf with BASTA® solution and observing sensitivity to the herbicide. The *grp8-5* allele harbored a 1 bp (A) insertion after position +27 and was created with sgRNA #7. The *grp8-11* allele harbored an insertion of 15 bp starting at position -5 and a following deletion of 334 bp, resulting in the loss of the complete first exon and part of the intron. The allele was obtained by combining sgRNAs #1 and #7.

### Generation of p35S::eYFP:GRP8 in Col-0 and Q plants

Plant expression vectors encoding fluorescently tagged GRP8 (eYFP-GRP8) under control of the Cauliflower Mosaci Virus 35S promoter were obtained by recombination between pENTR-GRP8 and empty pH7WGY2 ^51^ using LR clonase (ThermoFisherScientific). The resulting recombinant plasmids were then electroporated into *Agrobacterium tumefaciens* GV3101 pMP90. The floral dip method introduced the construct into Col-0 and Q plants. Hygromycin was used for transformant selection and monitoring the GFP signal in antibiotic-resistant seedlings.

### Generation of pGRP8::msfGFP:GRP8/*grp8-11* plants

The pGRP8::msfGFP:GRP8 expression cassette was designed to include 1.9 kb of the *GRP8* promoter and the native 5’UTR, followed by plant-optimized coding sequence of msfGFP (SuperfolderGFP with A203K mutation for monomerization), the genomic *GRP8* sequence harboring the intron, and the *GRP8* 3’UTR based on the previously published data ^21^. The entire cassette was ordered as a synthetic gene flanked by attB1 and attB2 sites (ThermoFisherScientific). The obtained plasmid was recombined with pDONR221 using BP clonase (ThermoFisherScientific), yielding pENTR221-pGRP8::msfGFP-GRP8 construct that was further recombined with pGWB601 to obtain the final construct in *E. coli/A. tumefaciens* shuttle vector. Final plasmid pGWB601-pGRP8::msfGFP-GRP8 was electroporated into *A. tumefaciens* GV3101 pMP90. The construct was introduced into *grp8-11* by the floral dip method. BASTA was used for transformant selection together with monitoring of GFP signal in BASTA-resistant seedlings.

### Expression and purification of recombinant proteins

A list of all constructs prepared in this study is provided in Supplementary Table 1.

### Expression and purification of Glutathione-*S-*transferase (GST) fusion proteins

Recombinant SnRK2.4, SnRK2.5, and SnRK2.10 kinases were prepared as described previously ^45^.

Cloning, expression, and purification of GST-GRP8 fusion proteins were performed essentially as described in ^18^, but instead of *ERD10*/*ERD14* cDNA, *GRP8* cDNA was PCR-amplified using specific primers listed in Supplementary Table 2.

### Expression and purification of msfGFP-GRP8 fusion proteins

To express msfGFP-GRP8 fusion proteins, the coding sequence of *GRP8* or its point mutants was amplified with primers bearing Bsp1407I (FW) and XhoI (RV) restriction sites (listed in Supplementary Table 2). The pET His_6_ MBP prescission LIC cloning vector (a gift from Scott Gradia; Addgene plasmid #29721; http://n2t.net/addgene:29721; RRID: Addgene_29721) was modified by insertion of msfGFP coding sequence from the pET MBP GFP LIC cloning vector (a gift from Scott Gradia; Addgene plasmid # 29750; http://n2t.net/addgene:29750; RRID: Addgene_29750) between SspI and XhoI sites and named pAMK2P. Purified PCR products and pAMK2P were digested with Bsp1407I and XhoI (ThermoFisherScientific), purified, and ligated. Positive clones were checked by Sanger sequencing. For protein expression, plasmids were transformed to *E. coli* BL21(DE3) pRARE and expressed overnight in 500 mL of ZYM-5052 autoinduction medium described by ^52^.

For purification of recombinant His_6_-MBP-msfGFP-GRP8 proteins, the bacteria were pelleted and resuspended in 40 mL His Wash Buffer (50 mM Tris/Tris-HCl, 500 mM NaCl, pH 8.0) supplemented with 1 mM PMSF, 1X Complete Inhibitor Cocktail EDTA free (Roche) and 250 μg/mL lysozyme. After 15 min of rotation at 4°C, Triton X-100 was added to a final concentration of 1% together with 1 μL of Viscolase (A&A Biotechnology), and the rotation was continued for another 15 min at 4°C. The lysates were sonicated for 5 min on ice and subsequently centrifuged for 20 min at 20,000 x g. Clarified lysate was supplemented with imidazole to 10 mM, filtered through a 0.45 μm cellulose acetate filter and subjected to FPLC purification on a 5 mL HisTrap HP column (Cytiva). The column was washed with 4% His Elution Buffer (50 mM Tris/Tris-HCl, 500 mM NaCl, 500 mM imidazole, pH 8.0) and eluted with 30% His Elution Buffer in His Wash Buffer. Fractions containing purified His_6_-msfGFP-GRP8 protein were pooled, and an appropriate amount of His-GST-tagged 3C HRV protease was added (1 μg of the protease per 1 mg of recombinant protein). The cleavage mixture was dialyzed overnight against 5 L of TBS (50 mM Tris/Tris-HCl, 150 mM NaCl, pH 7.5) at 4°C. Cleaved msfGFP-GRP8 was separated from His-MBP by FPLC using the 5 mL HisTrap HP column again, and flowthrough containing msfGFP-GRP8 was collected. Subsequently, msfGFP-GRP8 was purified on a 5 mL HiTrap Heparin HP column (Cytiva). Protein was loaded in 25 mM Tris/Tris-HCl with 25 mM NaCl and eluted with a linear 5-20% gradient of 25 mM Tris/Tris-HCl with 2M NaCl. Fractions were analyzed by SDS-PAGE, and protein bands were visualized by 2,2,2-trichloroethanol staining at each purification step. Fractions containing pure protein were pooled for the next chromatographic step. After the final purification on the Heparin column, fractions containing pure protein were concentrated by ultrafiltration with a simultaneous buffer exchange to 100 mM Tris/Tris-HCl, 300 mM NaCl, pH 7.5. The solution was mixed 1:1 with glycerol and stored at -20°C. For use as a control, unfused msfGFP expressed from empty pAMK2P was purified using the same method except for the Heparin chromatography. All chromatographic steps were performed at 4°C.

### Expression and purification of untagged GRP8 variants for HDX experiments

The coding sequence of GRP8 or its variant was amplified with specific primers bearing BamHI and XhoI sites (listed in Supplementary Table 2). The pCIOX vector (a gift from Andrea Mattevi; Addgene plasmid #51300; http://n2t.net/addgene:51300; RRID: Addgene_51300) and purified PCR product were cleaved with BamHI and XhoI FastDigest enzymes (ThermoFisherScientific), ligated and transformed into *E. coli* DH10b. Plasmids from positive clones were verified by sequencing.

The recombinant pCIOX-GRP8 plasmid was transformed to *E. coli* Rosetta (DE3) and cultured in 500 mL LB for protein expression. The culture was induced with 0.5 mM IPTG when the OD_600_ reached about 0.8 and then incubated at 18°C overnight. Bacteria were harvested by centrifugation, and pellets were stored at - 20°C. His_8_-SUMO-GRP8 fusion proteins were purified essentially as described above for msfGFP-GRP8. After the first affinity step, pooled fractions were cleaved using His_6_-SenP2 (SUMO protease, BPSBioscience) and rechromatographed on the HisTrap HP column to obtain tag-free GRP8 variants. Subsequently, proteins were purified on the HiTrap Heparin column as described above, yielding pure untagged proteins that were concentrated and stored as described above.

### *In vitro* phosphorylation of recombinant GRP8 and identification of phosphorylation sites

*In vitro* phosphorylation of recombinant GST-GRP8 WT protein and the GST-GRP8^S27A^ variant using γ-[^32^P] ATP (Hartmann Analytic) was performed essentially as described by ^18^, but instead of ERD10/ERD14, 5 µg of recombinant GST-GRP8 or GST-GRP8^S27A^, or 4 µg of Myelin Basic Protein (MBP) as a kinase activity control were used.

For the LC/MS analysis, the phosphorylation was performed as above but without γ-[^32^P] ATP. The reaction was stopped by precipitation of proteins with chloroform/methanol.

### Immunoblotting analysis

Plant material (about 50 mg of roots per sample) was ground in liquid nitrogen, 2X Laemli sample buffer was added to powdered samples in the ratio of 2:1 (v:w), and after boiling (95°C for 5 min), 20 µL samples were separated by 4-15% SDS-PAGE (Mini-PROTEAN^®^ TGX™ Precast Protein Gels, BioRad). A mixture of recombinant proteins was used as a standard: GFP-GRP8 (66 ng per lane) + GRP8 (58 ng per lane). Separated proteins were transferred to the PVDF membrane (Amersham^TM^ Hybond^TM^) by electroblotting and visualized by staining with 2% Ponceau S in 3% TCA.

For immunodetection, rabbit anti-GRP8 antibodies ^27^ diluted 1:5000 in TBST with 5% milk were used. After a 1 h incubation in RT, the membranes were extensively rinsed with TBST and incubated for 1 h with goat anti-rabbit horseradish peroxidase (HRP)-conjugated secondary antibodies (Agrisera) diluted 1:25,000 in TBST with 3% milk. For HRP detection, Pierce ECL Western reagent was used according to the manufacturer’s protocol.

### Site-directed mutagenesis

Site-directed mutagenesis was performed using Quick Change II Site–Directed Mutagenesis Kit (Agilent) and primers listed in Supplementary Table 2. The mutated protein-coding sequences were verified by sequencing, and the plasmids were transformed into an appropriate *E. coli* strain for expression or into *Agrobacterium tumefaciens* strain GV3101 pMP90 for transient expression experiments and generation of transgenic plants.

### Transient expression in *Nicotiana benthamiana*

Constructs for intracellular localization of proteins were prepared using the Gateway^®^ Cloning System and primers listed in Supplementary Table 2. *GRP8* cDNA was PCR-amplified and cloned into the pENTR-MCS2 vector, which was obtained by insertion of 58 bp oligonucleotide containing EcoRI, BshTI, SalI, XhoI SpeI, BamHI, and KpnI sites into pENTR^™^/D-TOPO^™^ vector cleaved with NotI and PteI. Mutated GRP8 variants were generated as described above. Then, the required cDNA was recombined into pSITE-2CA vector by Gateway LR reaction (ThermoFisherScientific) and transformed into the *A. tumefaciens* strain GV3101 pMP90.

For transient expression of the constructs in *N. benthamiana* leaves, fresh overnight cultures of *A. tumefaciens* containing appropriate binary plasmids were spun down and washed twice with sterile water. For localization experiments, the bacteria were resuspended in sterile water to a final density of 3×10^8^ cfu/mL (OD_600_ ∼ 0.3). For co-localization assays, appropriate bacterial suspensions at 6×10^8^ cfu/mL were combined in a 1:1 ratio before infiltration. Leaves of 4–5-week-old *N. benthamiana* plants were infiltrated with the bacterial suspension using a needleless syringe. The leaves were harvested and analyzed under a confocal microscope 2 days after agroinfiltration.

### Transient expression of RFP-fusion proteins in *A. thaliana*

Plants expressing eYFP-GRP8 (eYFP-GRP8-WT) or *grp8.11* were grown in a growth chamber under short-day conditions (8 h light/16 h dark photoperiod) at 22°C/19°C in soil. To introduce RFP-UBP1 or RPF-DCP1 constructs into the leaves of eYFP-GRP8-WT or *grp8-11* plants, *A. tumefaciens* containing appropriate binary plasmids were cultured as described by ^53^. Briefly, an overnight culture was spun down and resuspended in the induction medium (10.5 g/L K_2_HPO_4_, 4.5 g/L KH_2_PO_4_, 1 g/L (NH_4_)_2_SO_4_, 0.5 g/L Na_3_Citrate, 1 g/L glucose, 1 g/L fructose, 4 g/L glycerol, 1 mM MgSO_4_, 10 mM MES, pH 5.6) supplemented with 100 μM acetosyringone and incubated at 28°C for 5-6 hours with shaking. A calculated volume of the suspension was spun down again and resuspended in an appropriate volume of infiltration medium (10 mM MES pH 5.6, 10 mM MgSO_4_) supplemented with 200 μM acetosyringone to obtain a final density of 5x10^8^ CFU/ml (OD_600_=0.5). The abaxial side of the leaf was then infiltrated with the suspension as above. Plants were grown as above for 3 days, and leaves were analyzed using a confocal microscope.

### Confocal laser scanning microscopy

Microscopy was performed on the confocal laser scanning system built on an inverted microscope (EZ-C1 v.3.91, TE2000E, Nikon Instruments B.V. Europe, Amsterdam, The Netherlands). GFP and YFP were excited at 488 nm laser (Sapphire 488-20, 20 mW; Coherent, USA). Their emissions were detected by a 515/30 nm single-band bandpass filter (BrightLine, Semrock) and 610 long-pass filter (Chroma), respectively. RFP was excited at 543 nm laser (Green He-Ne Laser, 1.0 mW; Melles Griot, USA), and the fluorescence was collected by a 607/70 nm single-band bandpass filter (BrightLine, Semrock). Fluorescence signals were gathered with an oil-immersion 60x objective (N.A. 1.4, CFI Plan-Apochromat, Nikon). To avoid crosstalk and photobleaching, for every set of experiments, low laser power, and single-channel acquisition were defined, pinhole and exposure time were also optimized. At least ten images were collected from three biological replicates for each treatment.

In FRAP experiments, the following settings of the 488 nm laser were used: 0.1% power for pre-bleaching and recovery imaging and 100% power for bleaching. Sequences were created from 10 pre-bleach images, one scan of bleaching in the ROI circular area 2 μm^2^, and 100 recovery images. Images were collected at 256 pixels^2^ with a scan speed of 0.590 sec/frame. We tested that the energy of the 488 nm laser used to record the data after bleaching was weak enough that there was no signal drop-off in the unbleached ROIs. For analysis of the FRAP, the data were processed in FIJI/ImageJ (https://imagej.net/software/fiji/) software and calculated accordingly to plugins from Stowers Institute (https://research.stowers.org/imagejplugins/ImageJ_tutorial2.html). At least ten measurements were collected from several seedlings, and a representation was selected for calculations. All fluorescent images were processed in EZ-C1 Viewer and FIJI/ImageJ, and figures were made in FigureJ (https://imagej.net/plugins/figurej).

### Molecular modeling

The structure of the N-terminal domain of GRP8 was modeled by homology using the Yasara Structure package (www.yasara.org). Eight template structures were automatically selected in the Protein Data Bank based on sequence similarity and structure quality (4C7Q, 2MPU, 3S7R, 5IM0, 5TBX, 7JLY, 2RRB, and 6Q2I). Up to five alternative alignments to the target sequence were tested for each template, with up to 50 alternative conformations tested for each modified loop. The resulting models were individually evaluated for structural quality (dihedral angles, backbone, and side chain packing). Those with the highest scores were then used to create a hybrid model built from the best fragments identified among the single-template models. The resulting model was based on *Homo sapiens* cold-inducible RNA-binding protein (pdb record 5TBX) supplemented locally with *Nicotiana tabacum* RNA-binding glycine-rich protein (4C7Q). The sidechain rotamers were then tuned with ten rounds of the “repairPDB” procedure implemented in the FoldX5 package (DOI: 10.1093/bioinformatics/btz184). The effects of S27 phosphorylation and the S27D, S27E, and S27A, replacements on the stability of the modeled domain were then assessed with the FoldX5 “PositionScan” procedure.

The relative orientation of the N-terminal GRP8 domains in the dimer was adapted from the crystal structure of the *Trypanosoma brucei* RRM domain in complex with RNA (6E4P). The oligonucleotide location was then taken from the solution structure of *Homo sapiens* Polypyrimidine Tract Binding protein RBD1 complexed with CUCUCU RNA (2AD9). The final structure of the RNA-bound dimer was tuned with the Simulated Annealing procedure implemented in Yasara Structure. The change in stability of the complex upon introduction of amino acid replacement was assessed with the FoldX5 “AnalyseComplex” procedure.

### GRP8 protein sequence analysis

PrLD predictions for GRP8 were performed using PLAAC (http://plaac.wi.mit.edu/) using default settings and full-length protein sequence.

### Hydrogen-Deuterium Exchange Monitored by Mass Spectrometry (HDX-MS)

The analysis was performed for GRP8^WT^ and GRP8^S27D^. HDX exchange incubation was performed for 10 s, 1 min, 5 min, 30 min, and 2.5 h at room temperature with 5 μL of protein stock (50 μM) added to 45 μL of deuterated buffer (50 mM Tris-HCl pH 7.0, 300 mM NaCl). The H/D exchange was quenched by transferring the protein solution to a precooled tube (on ice) containing 10 μL of quenching buffer (2 M glycine in 99.99% D_2_O, pD 2.3). After quenching, the samples were immediately flash-frozen in liquid nitrogen and kept at -80°C until LC-MS analysis, carried out within 2-3 days.

HDX samples were thawed directly before measurement and injected manually into a nano ACQUITY UPLC system equipped with HDX-MS Manager (Waters). Proteins were digested on a 2.1 mm × 20 mm column with immobilized pepsin (AffiPro) for 1.5 min at 20°C and eluted with 0.07% formic acid in water at a 200 μL/min flow rate. Digested peptides were passed directly onto an ACQUITY BEH C18 VanGuard pre-column followed by a reversed-phase ACQUITY UPLC BEH C18 column (Waters) using a 10–35% gradient of acetonitrile in 0.01% formic acid at a flow rate of 90 μL/min at 0.5°C. The eluate was subjected to MS analysis on a SYNAPTG2 instrument (Waters) with the following parameters: ESI — positive mode; capillary voltage — 3 kV; sampling cone voltage — 35 V; extraction cone voltage — 3 V; source temperature — 80°C; desolvation temperature — 175°C; and desolvation gas flow 800 L/h.

Two control experiments were conducted to assess the minimum and maximum H-D exchange levels for each peptide. For the minimum exchange value (Mmin), 10 µL of quench buffer was mixed with 45 µL of the D_2_O exchange buffer prior to the addition of 5 µL of protein stock followed by the LC-MS procedure. For the maximum exchange value (Mmax), the deuteration reaction was conducted for 24 hours and then quenched with a quench buffer kept on ice.

Peptide lists for each GRP8 variant were obtained for protein samples subjected to a mock-HDX procedure (H_2_O used instead of D_2_O). The peptides were identified using ProteinLynx Global Server Software (Waters). The peptide lists obtained for non-deuterated proteins were used to analyze the exchange data using DynamX 3.0 (Waters) software. First, the peptide lists were filtered by the following criteria: minimum intensity — 3000 and minimal product per amino acid — 0.3. All raw files were processed and analyzed in DynamX 3.0 software. All DynamX assignments were inspected manually.

The fractional percentage of deuteration D [%] for each peptide was calculated using an in-house HaDeX data analysis software (Puchała et al., 2020) from exported DynamX 3.0 data, using the formula taking into account the minimal and maximal exchange of a given peptide and mean values of exchange calculated from at least 3 replicates:

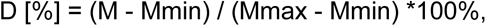

where M – means the centroid mass of a given peptide after deuterium uptake, Mmin – means the centroid mass of the peptide with a minimal exchange, and Mmax – means the centroid mass of the peptide with a maximal exchange. Kinetic plots of the deuterium uptake for the GRP8 peptides at different incubation times were obtained using Origin 9.1.0 (OriginLab Corp.) software. To overlay the fractional deuteration levels on the GRP8 structural model, UCSF Chimera software (developed by the Resource for Biocomputing, Visualization, and Informatics at the University of California, San Francisco, with support from NIH P41-GM103311) ^54^ was used.

### RNA electromobility shift assay (REMSA)

For RNA binding experiments, 250 fmol of Cy5-labelled RNA (Supplementary Table 2) was incubated with 0-1000 fmol of suitable GST-GRP8 fusion protein variant in a total volume of 20 μL of 1X EMSA buffer (20 mM HEPES/HEPES-Na, 100 mM NaCl, 0.01% NP-40) containing 2 μg of yeast tRNA (Sigma-Aldrich) and 10 U of RiboLock (ThermoFisherScientific) for 30 min at 22°C. Then 2.2 μL of 10X EMSA loading buffer (50% glycerol, 0.01% Orange G) was added, and samples were subjected to PAGE electrophoresis in 6% gel (75:1 crosslink ratio) with 10% glycerol in 0.5X TBE. Gels were scanned using a Typhoon FLA9000 laser imager; densitometry of the bands was performed using ImageJ software.

### *In vitro* liquid-liquid phase separation (LLPS) assay

The phase separation assay was performed in 384-well clear F-bottom black plates (Greiner Bio-One #781986). Protein of interest (suitable msfGFP-GRP8 fusion protein variant or msfGFP alone) at 2X final concentration required was mixed 1:1 with an appropriate 2X concentrated LLPS buffer consisting of 25 mM HEPES:HEPES-Na pH 7.5 and KCl and PEG3350 at various concentrations (in some variants, total Arabidopsis RNA at indicated concentrations was added in the presence or absence of PEG), shaken for 30 s and briefly spun down at 500 RCF. Unless indicated otherwise, the optimized, 2X concentrated buffer was used of the following composition: 25 mM HEPES:HEPES-Na pH 7.5, 250 mM KCl, 24% PEG 3350. LLPS was imaged after 30 min of incubation at RT using an Olympus IX81 wide-field fluorescence microscope equipped with a ScanR modular imaging platform (Olympus) and a UPlanSApo 20×/0.75 objective. ScanR Acquisition software (version 3.0.0) was used for microscope control. Protein droplets were imaged using GFP excitation and emission wavelength. Image analysis was performed in ScanR Analysis software (version 3.0.0). Images of the GFP channel were subjected to rolling-ball background subtraction, and segmentation was performed using the Edge algorithm. Each identified object was assumed to be a protein droplet.

Further numerical data processing was performed using the R environment. For each experimental condition, the total number of droplets was calculated, and conditions in which the number of droplets was less than 0.5% of the maximum number were considered unfavorable to LLPS. The phase-separated fraction was calculated as follows: (Int_droplet_ * Area_droplet_ / (Int_image_ * Area_image_) where Int_droplet_ is the average of mean droplet fluorescence intensities, Area_droplet_ is the sum of all droplet areas, Int_image_ is the mean fluorescence intensity of the image, and Area_image_ is the size of the image ^35^. Statistical significance was calculated using an appropriate omnibus test. All data were checked beforehand for normality using the Shapiro-Wilk test and the equality of variances using Bartlett’s test.

For montages, representative images obtained by the abovementioned imaging procedure were selected. Post-acquisition image processing, performed in Fiji/ImageJ software (version 1.53f51), included cropping to provide a more accurate visualization of protein droplets and rolling-ball background subtraction.

### Glutaraldehyde crosslinking and SDS-PAGE

Recombinant GFP-GRP8 proteins at the indicated concentrations were crosslinked for 30 min at 22°C in buffer containing 25 mM HEPES:HEPES-Na, 125 mM NaCl pH 7.5, with 10 mM glutaraldehyde. The reaction was terminated by adding 5X concentrated SDS-PAGE reducing sample buffer and boiling at 95°C for 5 min. Three µg of protein was run on a 4-20% gradient SDS-PAGE gel (Bio-RAD), and protein bands were visualized by Coomassie Blue staining.

### Dynamic light scattering measurements

Dynamic light scattering of GFP-GRP8 and its variants (5 μM) in 25 mM HEPES:HEPES-Na pH 7.5 with 125 mM KCl was determined at 25°C with a DynaPro NanoStar (Wyatt), using disposable microcuvettes (Eppendorf). Ten 10-s autocorrelations were recorded for each sample, and each measurement was repeated at least five times. The hydrodynamic radius and apparent molecular weight estimates were analyzed with the manufacturer’s software.

## ACKNOWLEDGEMENTS

We are grateful to Dr. Jan Fronk for the critical reading of the manuscript. We want to thank Dr. Marcin Nowotny and Dr. Mariusz Czarnocki-Cieciura for helping us set up the DLS experiment, Lilia Zukowa for performing HDX-MS experiments, Marta Piecho-Kabacik for helping us grow plants, Dr. Artur Jarmołowski and Dr. Zofia Kulińska-Szweykowska and all laboratory members for stimulating discussions. This work was supported by the National Science Centre grants: 2016/23/B/NZ3/03182 to G.D. and 2019/33/N/NZ3/02027 to A.K., German Research Foundation grant STA 653/14-1 to D.S.

---

**Extended data Figure 1.**
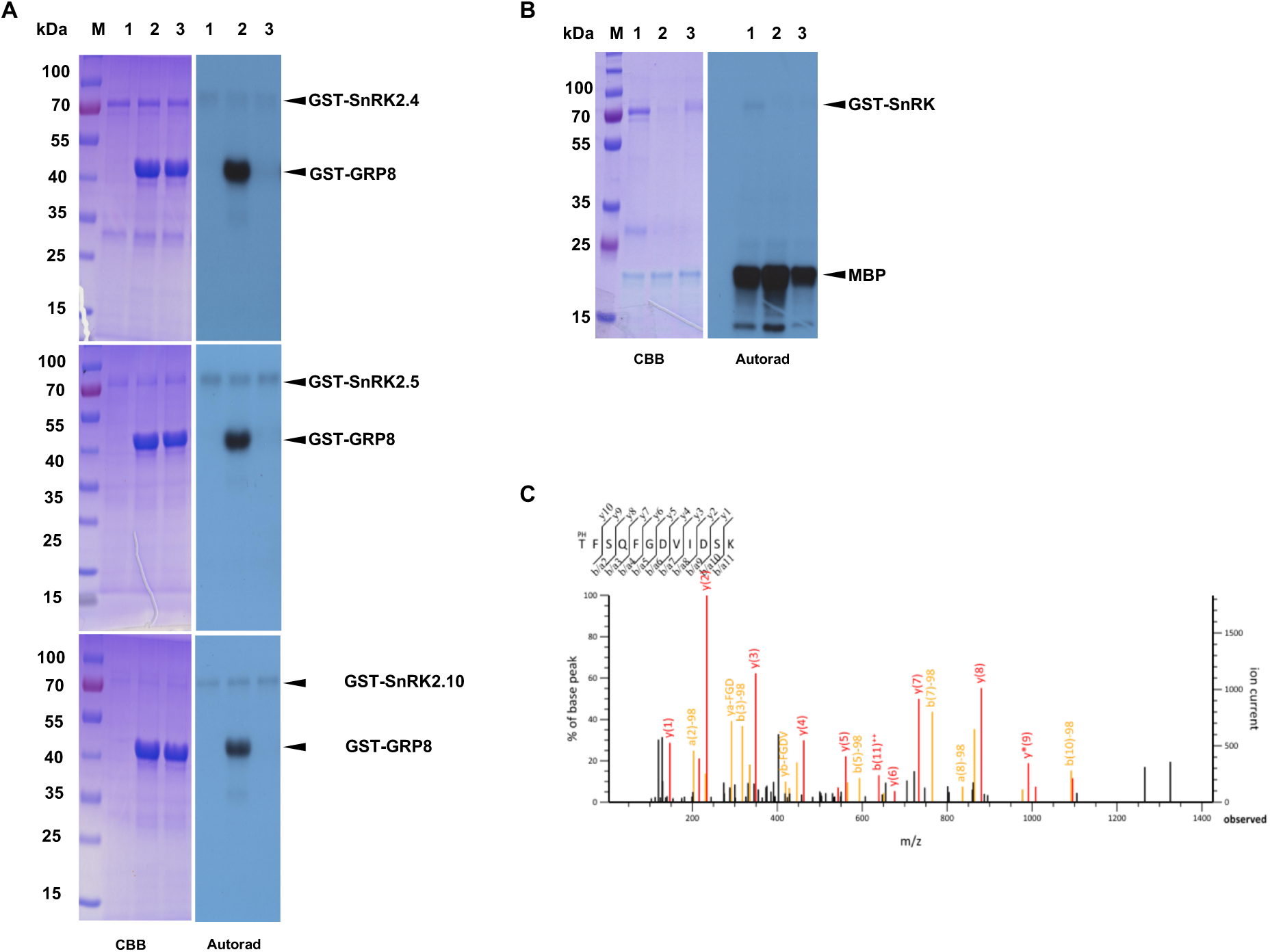
(A) GST-GRP8 *in vitro* phosphorylation by purified GST-SnRK2.4, GST-SnRK2.5, and GST-SnRK2.10; CBB –Coomassie Blue staining, Autorad – visualization of ^32^P signal of phosphorylated proteins; 1 – kinase only control, 2 – kinase + GST-GRP8^WT^, 3-kinase + GST-GRP8^S27A^ (B) Activity control of the kinases used for GST-GRP8 phosphorylations presented in (A) using myelin basic protein (MBP) as a substrate; 1 –GST-SnRK2.4, 2 – GST-SnRK2.5, and 3 – GST-SnRK2.10; CBB –Coomassie Blue staining, Autorad – visualization of ^32^P signal of phosphorylated proteins; (C) Fragmentation mass spectrum of GRP8 phospho-peptide 25-36 identified upon *in vitro* phosphorylation of GST-GRP8 by GST-SnRK2.10 Results of one of three independent experiments showing similar results are shown.

**Extended data Figure 2.**
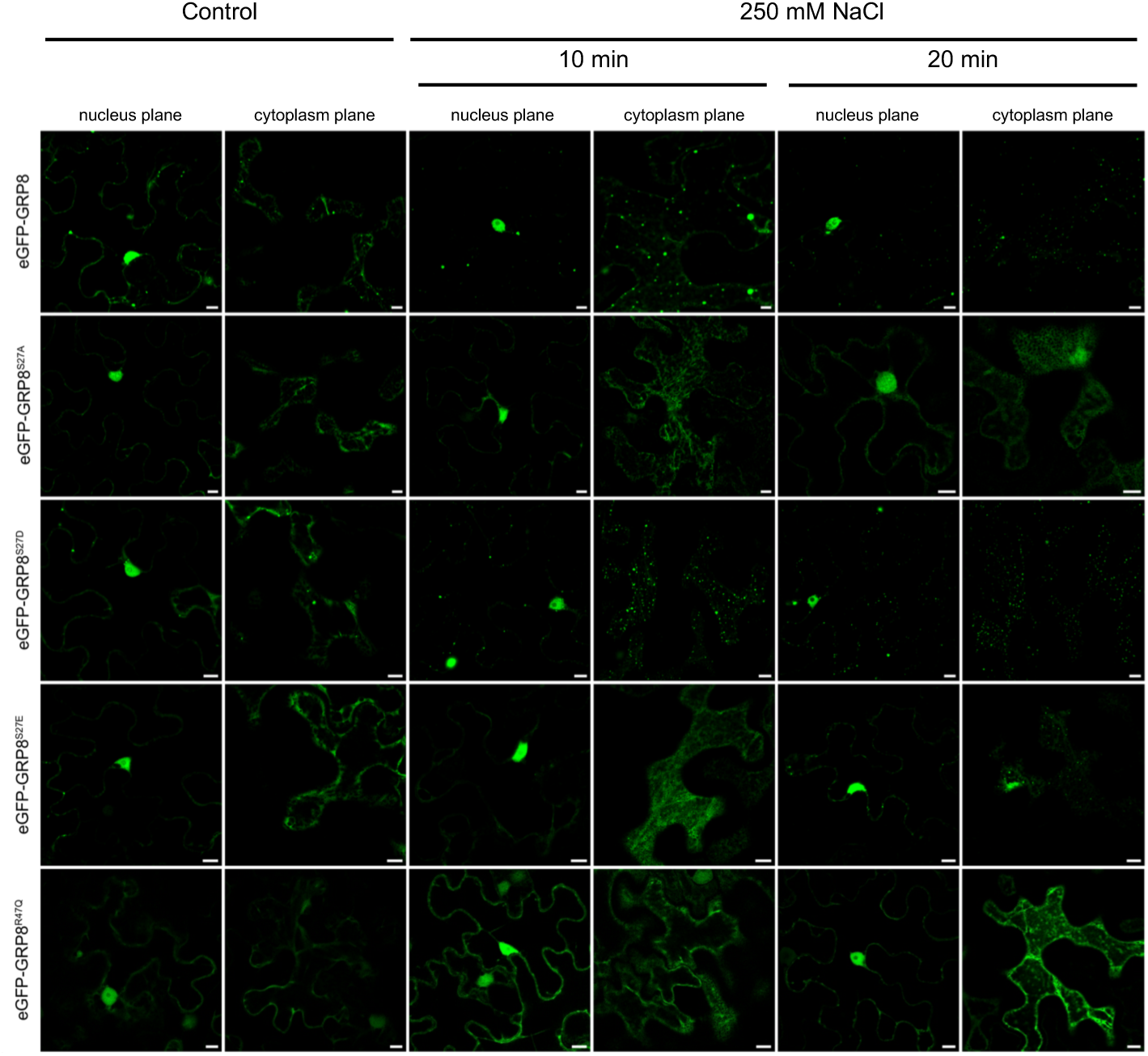
Confocal images of eGFP-GRP8 and its variants in transiently transformed *N. benthamiana* epidermal cells exposed or not (control) to salt stress (250 mM NaCl) including nucleus plane images. Scale bar represents 10 µm

**Extended data Figure 3.**
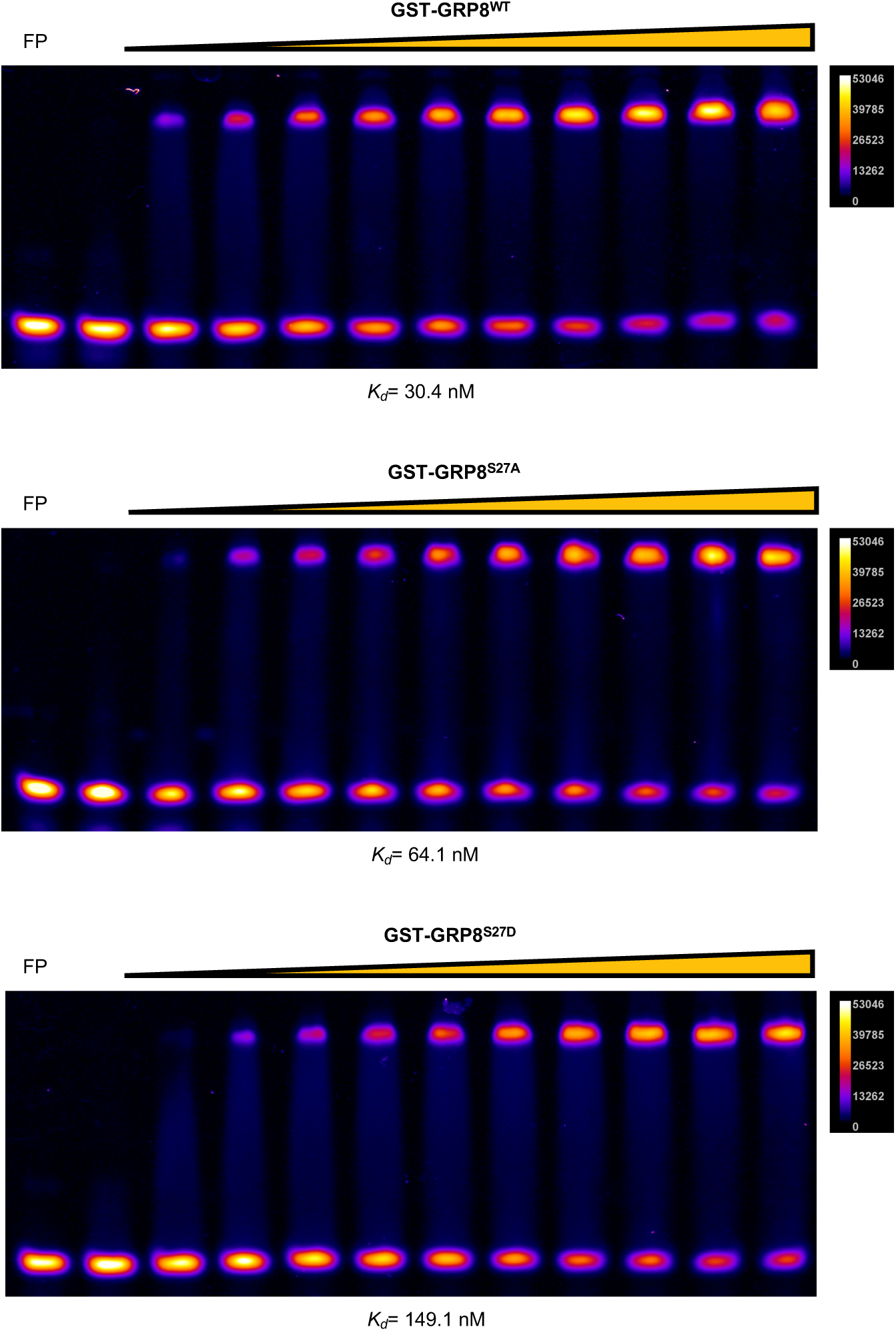
GST-GRP8^WT^, GST-GRP8^S27A^, and GST-GRP8^S27D^ binding with 3’-Cy5 labeled RNA oligonucleotide GRP8_3’UTR_WT; binding constants were calculated based on densitometric analysis of images using ImageJ and fitting nonlinear regression line in GraphPad Prism. Pixel intensity is indicated on the color scale; FP stands for free probe without protein added. Results of one of three independent experiments showing similar results are shown.

**Extended data Figure 4.**
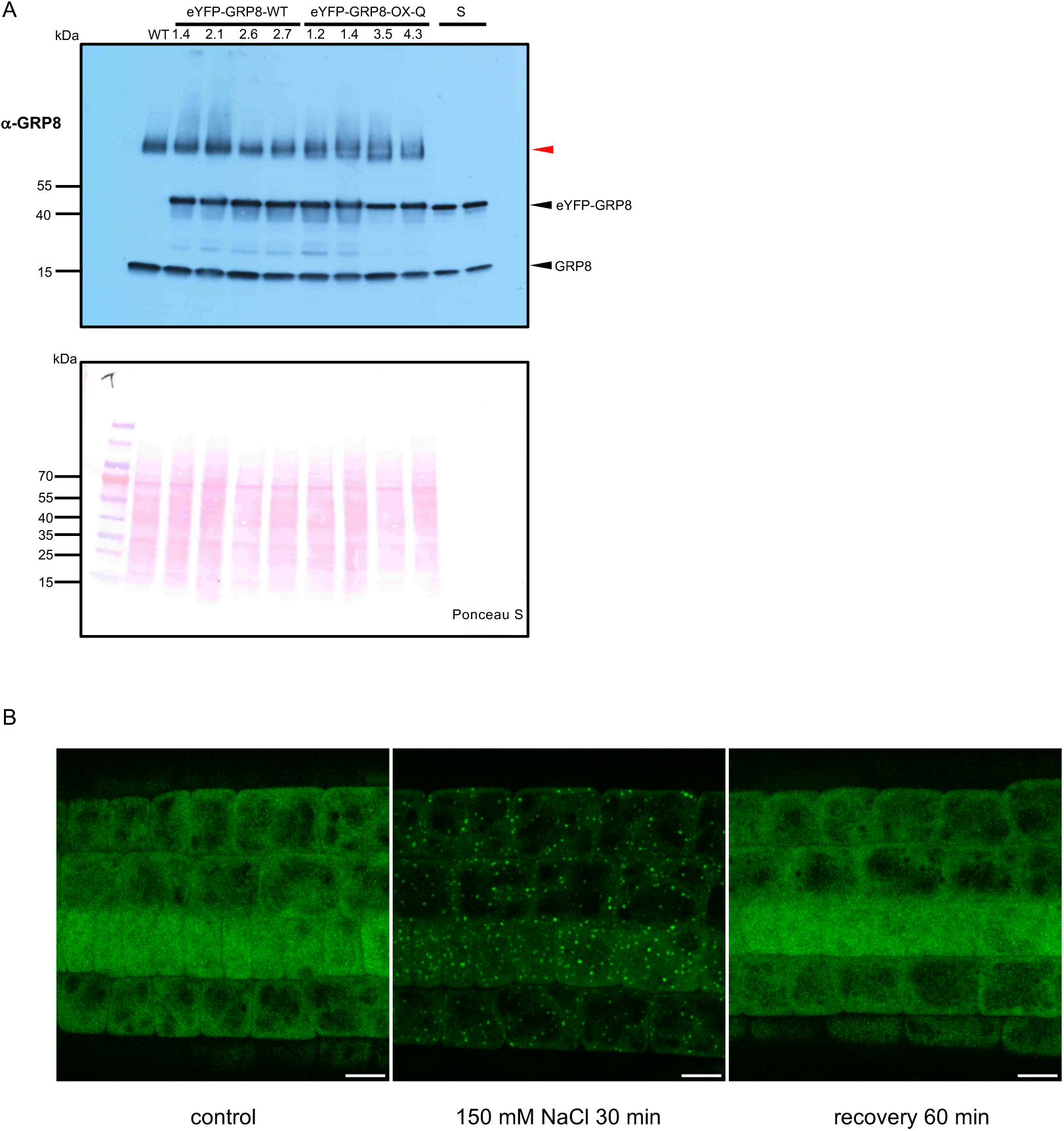
(A) Western blotting analysis of eYFP-GRP8 and native GRP8 levels in four independent lines of eYFP-GRP8-WT and eYFP-GRP8-Q plants using anti-GRP8 antibodies; S is a standard prepared by mixing purified recombinant msfGFP-GRP8 (66 ng per lane) and untagged GRP8 (58 ng per lane), Ponceau S staining of the membrane after transfer was used as a loading control. Red arrowhead indicates unspecific band. (B) Reversibility of GRP8 assembly in SGs in pGRP8::msfGFP-GRP8/*grp8-11* roots. Granules formed after 30 min of treatment with 150 mM NaCl disappeared after replacing the salt solution with growth medium for 1h. Results of one of three independent experiments showing similar results are shown. Scale bar = 10 μm.

**Extended data Figure 5.**
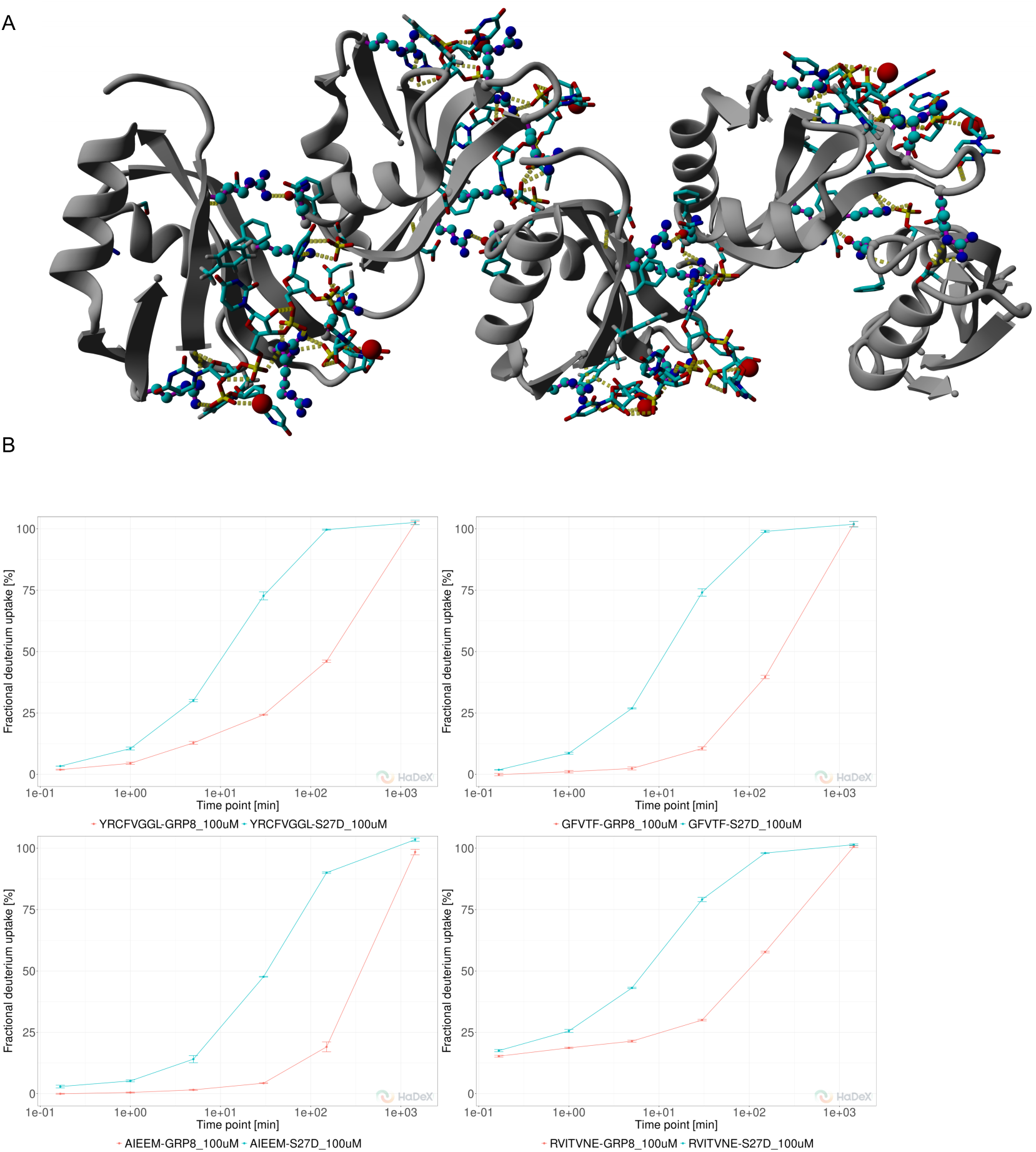
(A) In silico model of the GRP8 N-domain oligomer with RNA bound. (B) Deuterium uptake at different incubation times in GRP8^WT^ (red) and GRP8^S27D^ (blue) for four selected peptides characterized by the highest stability in WT and the largest difference in the deuterium uptake extent between WT and S27D: YRCFVGGL (upper left panel) covering region 6-13, GFVTF 50-54 (upper right panel), AIEEM 63-67 (left lower panel), RVITVNE 75-81 (right lower panel). The deuterium uptake mean value of the three replicate experiments is shown.

**Extended data Figure 6.**
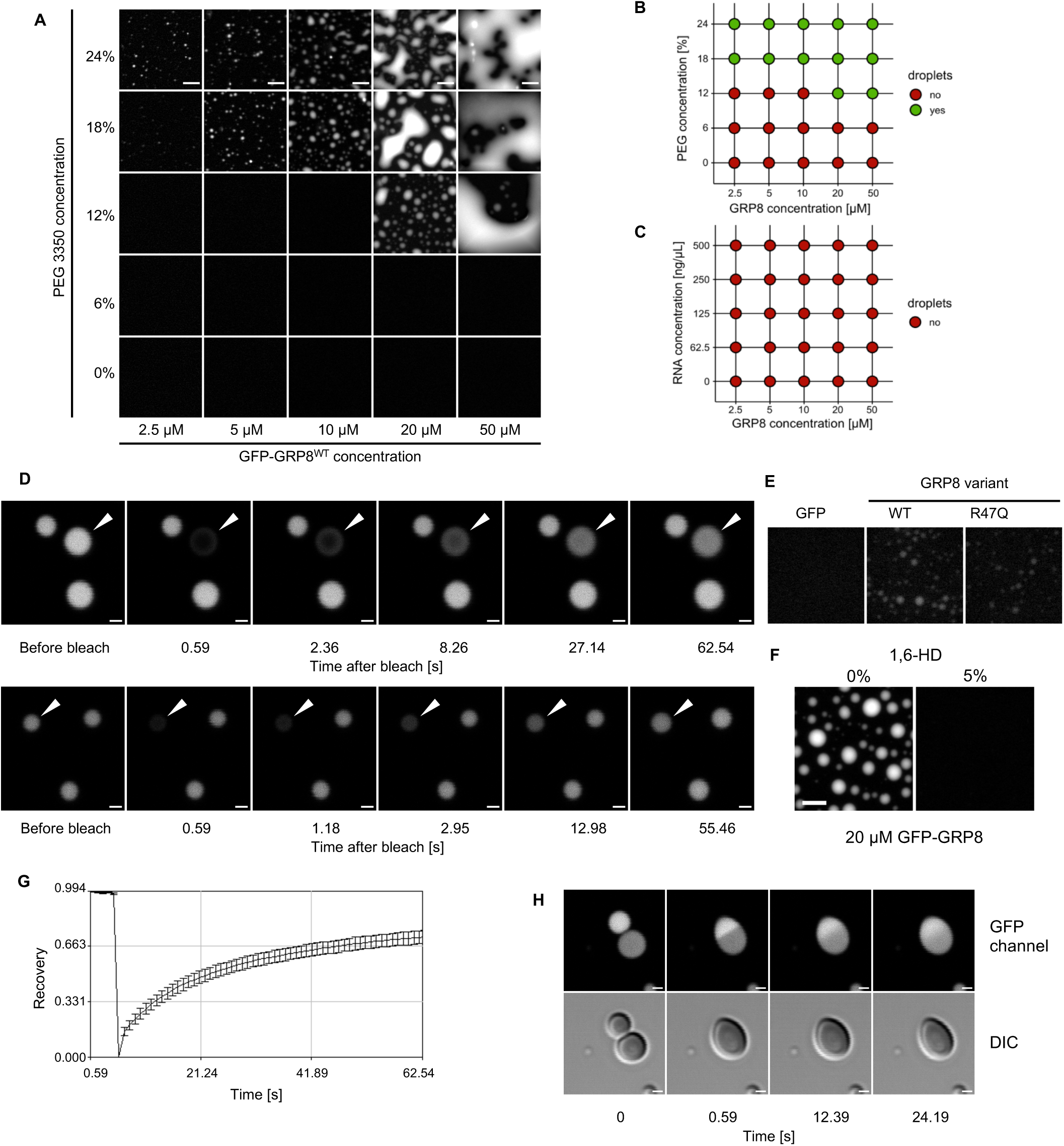
(A and B) Purified msfGFP-GRP8 phase separation in the presence of PEG 3350, Scale bar represents 10 µm; (C) *In vitro* LLPS of msfGFP-GRP8 in the presence of total Arabidopsis RNA without PEG; (D) FRAP assay on *in vitro* separated msf-GFP-GRP8 droplets, bleached droplet indicated by an arrowhead, Scale bar represents 1 µm; (E) *In vitro* LLPS of msfGFP-GRP8^R47Q^ under optimized assay conditions at 20 µM protein concentration; (F) Dissolution of droplets of msfGFP-GRP8 formed by LLPS *in vitro* by 1,6-hexanediol (1,6-HD); scale bar represents 10 µm (G) Time course of relative recovery after photobleaching of droplets of msfGFP-GRP8 formed by LLPS *in vitro s*hown in C). Data are mean +/-s.d. (n=7 independent experiments) (H) Fusion of droplets of msfGFP-GRP8. The larger droplet was photobleached before fusion to demonstrate the dynamic mixing of the two droplets upon coalescence. DIC = differential interference contrast, Scale bar represents 1 µm;

**Extended data Scheme 1.**
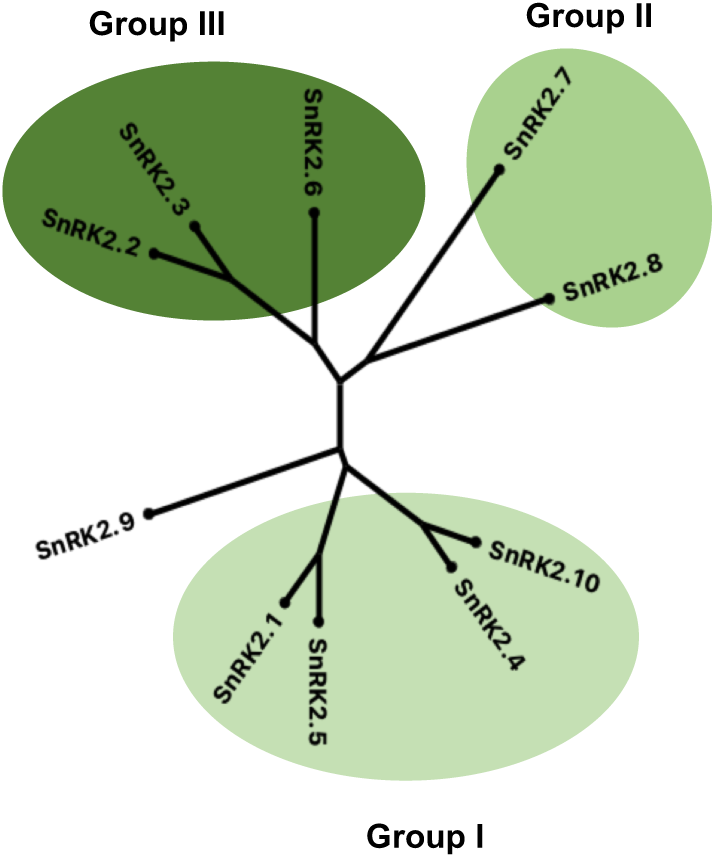
Phylogenetic tree of Arabidopsis SnRK2s. Group I consists of ABA-non-activated SnRK2s, group II - SnRK2s not or weakly activated in response to ABA, group III - SnRK2s strongly activated by ABA

**Extended Data Table 1.**
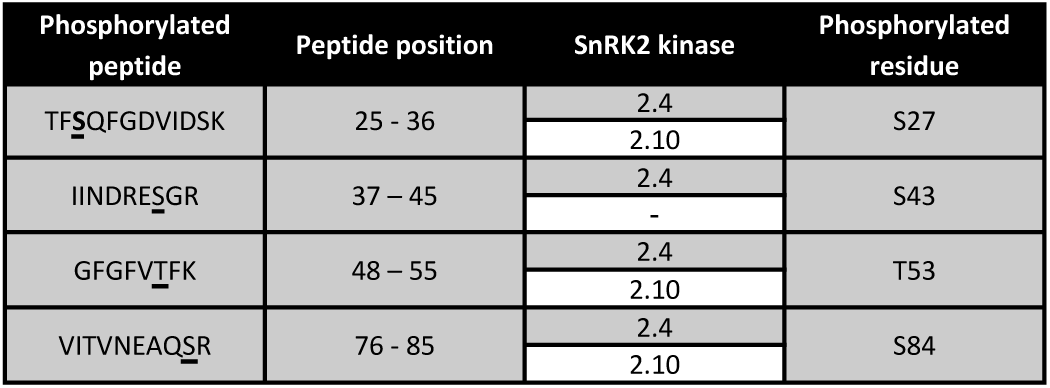
Phosphopeptides derived from GRP8 phosphorylated by SnRK2.10 and SnRK2.4 *in vitro.* Two independent experiments were performed for each kinase. The phosphorylated S27 residue identified in both experiments as phosphorylated by both kinases is bolded.

| Phosphorylated peptide | Peptide position | SnRK2 kinase | Phosphorylated residue |
| --- | --- | --- | --- |
| TF**S**QFGDVIDSK | 25 - 36 | 2.4 | S27 |
|  |  | 2.10 |  |
| IINDRE**S**GR | 37 – 45 | 2.4 | S43 |
|  |  | - |  |
| GFGFV**T**FK | 48 – 55 | 2.4 | T53 |
|  |  | 2.10 |  |
| VITVNEAQ**S**R | 76 - 85 | 2.4 | S84 |
|  |  | 2.10 |  |

