## Supplemental Table 1 and 2 for "Phosphorylation Promotes Liquid-Liquid Phase Separation of GRP8 and Its Assembly into Stress Granules Upon Salinity Stress in Arabidopsis"

**Supplementary Table 1: List of plasmids**

| Plasmid name | Description |
|---|---|
| pGEX-6P1-GRP8 | Protein expression for in vitro phosphorylation/EMSA |
| pGEX-6P1-GRP8$^{S27A}$ | Protein expression for in vitro phosphorylation/EMSA |
| pGEX-6P1-GRP8$^{S27D}$ | Protein expression for in vitro phosphorylation/EMSA |
| pAMK2P | Protein expression for in vitro LLPS, crosslinking and DLS |
| pAMK2P-GRP8 | Protein expression for in vitro LLPS, crosslinking and DLS |
| pAMK2P-GRP8$^{S27A}$ | Protein expression for in vitro LLPS, crosslinking and DLS |
| pAMK2P-GRP8$^{S27E}$ | Protein expression for in vitro LLPS, crosslinking and DLS |
| pAMK2P-GRP8$^{S27D}$ | Protein expression for in vitro LLPS, crosslinking and DLS |
| pAMK2P-GRP8$^{R47Q}$ | Protein expression for in vitro LLPS, crosslinking and DLS |
| pAMK2P-GRP8$^{R7A}$ | Protein expression for in vitro LLPS, crosslinking and DLS |
| pAMK2P-GRP8$^{Q28A}$ | Protein expression for in vitro LLPS, crosslinking and DLS |
| pAMK2P-GRP8$^{K36A}$ | Protein expression for in vitro LLPS, crosslinking and DLS |
| pAMK2P-GRP8$^{R41A}$ | Protein expression for in vitro LLPS, crosslinking and DLS |
| pAMK2P-C-GRP8 | Protein expression for in vitro LLPS |
| pAMK2P-N-GRP8 | Protein expression for in vitro LLPS |
| pCIOX-GRP8 | Protein expression for HDX-MS |
| pCIOX-GRP8$^{S27D}$ | Protein expression for HDX-MS |
| pENTR-MCS2-GRP8 | Entry clone for Gateway Recombination |
| pENTR-MCS2-GRP8$^{S27A}$ | Entry clone for Gateway Recombination |
| pENTR-MCS2-GRP8$^{S27D}$ | Entry clone for Gateway Recombination |
| pENTR-MCS2-GRP8$^{S27E}$ | Entry clone for Gateway Recombination |
| pENTR-MCS2-GRP8$^{R47Q}$ | Entry clone for Gateway Recombination |
| pENTR-MCS2-N-GRP8 | Entry clone for Gateway Recombination |
| pENTR-MCS2-C-GRP8 | Entry clone for Gateway Recombination |
| pSITE-2CA-GRP8 | Localisation and colocalisation experiments |
| pSITE-2CA-GRP8$^{S27A}$ | Localisation and colocalisation experiments |
| pSITE-2CA-GRP8$^{S27D}$ | Localisation and colocalisation experiments |
| pSITE-2CA-GRP8$^{S27E}$ | Localisation and colocalisation experiments |
| pSITE-2CA-GRP8$^{R47Q}$ | Localisation and colocalisation experiments |
| pSITE-2CA-N-GRP8 | Localisation and colocalisation experiments |
| pSITE-2CA-C-GRP8 | Localisation and colocalisation experiments |
| pGWB461-UBP1b | Co-localisation expereiments |
| pGWB461-RBP47b | Co-localisation expereiments |
| pGWB461-DCP1 | Co-localisation expereiments |
| pH7WGY2-GRP8 | Stable plant transformation, 35S driven expression |
| pMA-RQ-pGRP8::msfGFP-GRP8 | Syntetic expression casette |
| pENTR221-pGRP8::msfGFP-GRP8 | Entry clone with pGRP8 driven expression casette |
| pGWB601-pGRP8::msfGFP-GRP8 | Stable plant transformation, expression under pGRP8 native promoter |

**Supplementary Table 2: List of oligonucleotides**

**Primers for cloning**

| Name | Sequence | Comments |
|---|---|---|
| EcoRI-GRP8 FW | TTTGAATTCATGTCTGAAGTTGAGTACCGGTGCTTTG | GPR8 cloning into pGEX-6P1, pCIOX and pENTR-MCS2 |
| SalI-GRP8 RV | TTTGTCGACTTACCAGCCGCCACCACCG | |
| Bsp1407I-GRP8 FW | TATATGTACAAGATGTCTGAAGTTGAGTACCGGTGCTTTG | for cloning into pAMK2P downstream of GFP |
| XhoI-GRP8 RV | ATATCTCGAGTTACCAGCCGCCACCACCGCCTCCGTAAC | |

**Primers for CRISPR vectors**

| Name | Sequence | Comments |
|---|---|---|
| F_pEN_NotI | TTGCGGCCGCGCTTTTTTTCTTCTTCTTCGTTCATAC | |
| R_pEN_NotI | TTGCGGCCGCAAAAAAAAGCACCGACTCGGTG | |
| F_pEN_BamHI | TTGGATCCGCTTTTTTTCTTCTTCTTCGTTCATAC | |
| R_pEN_SpeI | TTACTAGTAAAAAAAGCACCGACTCGGTG | |

**Primers for mutagenesis**

| Name | Sequence | Comments |
|---|---|---|
| GRP8 S27A FW | CTTCAAAGGACGTTCGCACAGTTCGGCGACGTTATCG | |
| GRP8 S27A RV | CGATAACGTCGCCGAACTGTGCGAACGTCCTTTGAAG | |
| GRP8 S27E FW | CTTCAAAGGACGTTCGAACAGTTCGGCGACGTTATCG | |
| GRP8 S27E RV | CGATAACGTCGCCGAACTGTTCGAACGTCCTTTGAAG | |
| GRP8 S27D FW | GAAGATCTTCAAAGGACGTTCGACCAGTTCGGCGACGTTATCG | |
| GRP8 S27D RV | CGATAACGTCGCCGAACTGGTCGAACGTCCTTTGAAGATCTTC | |
| GRP8 R7A FW | GTTGAGTACGCGTGCTTTGTCGGCGGC | |
| GRP8 R7A RV | CCGACAAAGCACGCGTACTCAACTTCAG | |
| GRP8 K36A FW | CGATTCTGCGATCATTAACGACCGCG | |
| GRP8 K36A RV | TAATGATCGCAGAATCGATAACGTCG | |
| GRP8 Q28A FW | GTTCTCAGCGTTCGGCGACGTTATCG | |
| GRP8 Q28A RV | CGCCGAACGCTGAGAACGTCCTTTG | |
| GRP8 R41A FW | CATTAACGACGCCGAGAGTGGAAGATCAAGG | |
| GRP8 R41A RV | CCACTCTCGGCGTCGTTAATGATCTTAG | |
| GRP8 N-term FW INV | CTCGAGCACCACCACCACCACCACTG | GRP8 C-terminus deletion by PCR in pAMK2P-GFP-GRP8 |
| GRP8 N-term RV INV | TTAAGCCTCGTTCACGGTGATGACACG | |
| GRP8 C-term FW INV | CAGTCGAGAGGTAGCGGCGGT | GRP8 N-terminus deletion by PCR in pAMK2P-GFP-GRP8 |
| GFP-GRP8 C-term RV INV | CTTGTACAGCTCGTCCATGCCGAG | |

**RNA oligomers for EMSA**

| Name | Sequence | Comments |
|---|---|---|
| GRP8-3UTR-WT-Cy5 | GUUUUUGGUUUAGAUUUGGUUUUGUGU[Cyanine5] | for binding experiments |
